# Neurexin-3 defines synapse- and sex-dependent diversity of GABAergic inhibition in ventral subiculum

**DOI:** 10.1101/2021.04.22.440943

**Authors:** Emma E Boxer, Charlotte Seng, David Lukacsovich, JungMin Kim, Samantha Schwartz, Matthew J Kennedy, Csaba Földy, Jason Aoto

**Affiliations:** University of Colorado Anschutz, Department of Pharmacology, Aurora, CO USA 80045; Laboratory of Neural Connectivity, Brain Research Institute, Faculties of Medicine and Science, University of Zürich, Zürich, Switzerland 8057

## Abstract

Ventral subiculum (vSUB) is integral to the regulation of stress and reward, however the intrinsic connectivity and synaptic properties of the inhibitory local circuit are poorly understood. Neurexin-3 (Nrxn3) is highly expressed in hippocampal inhibitory neurons, but its function at inhibitory synapses has remained elusive. Using slice electrophysiology, imaging, and single-cell RNA sequencing, we identify multiple roles for Nrxn3 at GABAergic parvalbumin (PV) interneuron synapses made onto vSUB regular spiking (RS) and burst spiking (BS) principal neurons. Surprisingly, we found that intrinsic connectivity and synaptic function of Nrxn3 in vSUB are sexually dimorphic. We reveal that vSUB PVs make preferential contact with RS neurons in males, but BS neurons in females. Furthermore, we determined that despite comparable Nrxn3 isoform expression in male and female PV neurons, Nrxn3 maintains synapse density at PV-RS synapses in males, but suppresses presynaptic release at the same synapses in females.

**Highlights:**

- Overall inhibitory strength in ventral subiculum is cell-type specific
- PV circuits in ventral subiculum are organized sex-specifically
- Nrxn3 function in PV interneurons depends on postsynaptic cell identity
- Nrxn3 has distinct functions at PV-RS synapses in females compared to males

Graphical Abstract

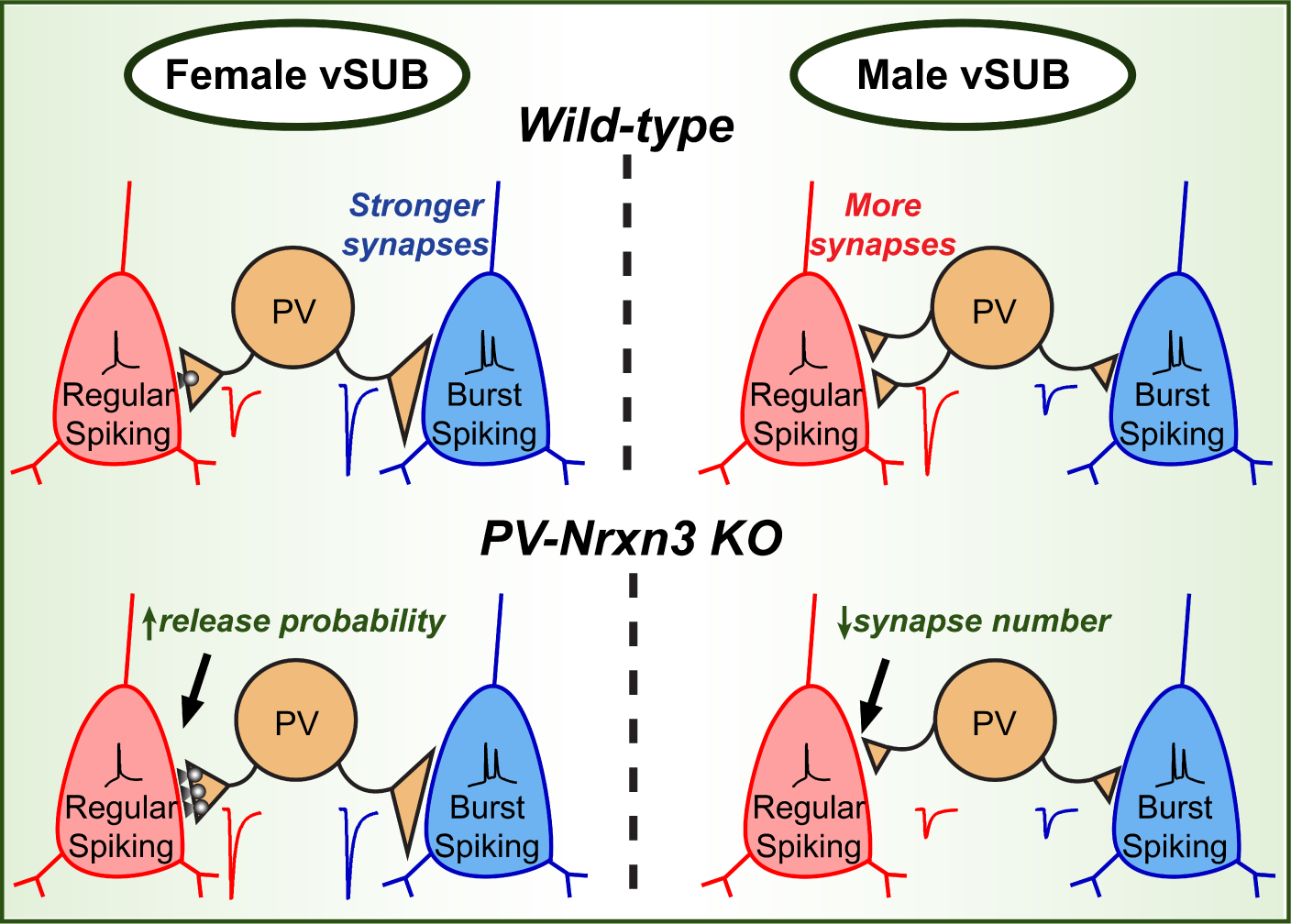

## Introduction

Precise innervation and synaptic transmission properties of local GABAergic interneurons with principal neurons are essential for faithful information processing and behavior, and disruption of inhibitory activity underlies many aspects of neuropsychiatric, neurodevelopmental and substance use disorders (SUDs). Synaptic cell adhesion molecules play a critical role in specifying and maintaining synaptic inhibition and mutations in genes that encode synaptic adhesion molecules are often linked to brain disorders. Neurexins (Nrxns) are a family of essential and evolutionarily conserved presynaptic cell adhesion molecules that engage in trans-synaptic interactions with an expanding number of postsynaptic ligands to control pre- and post-synaptic properties of synaptic transmission (Südhof, 2017). However, the cell-type and synapse-specific roles neurexins perform remain enigmatic. Further, stunning sex differences in the prevalence and symptomology of neuropsychiatric disorders such as schizophrenia (SCZ) and SUDs (Leung and Chue, 2003; Becker, 2016) raises the possibility that adhesion molecule usage in SCZ and SUD-relevant circuits is sexually dimorphic.

Neurexins are encoded by three genes (*NRXN1*-3), which make α, β, or the Nrxn1-specific γ isoforms from independent promoters, and are alternatively spliced at six conserved splice sites (SS1-6). While Nrxns1-3 exhibit high sequence homology, share common binding partners, and display some compensatory behavior (Aoto *et al*., 2015), recent evidence supports the hypothesis that individual neurexins play functionally distinct roles in synaptic transmission (Aoto *et al*., 2015; Dai *et al*., 2019). This is further supported by the highly variable expression patterns of neurexin isoforms and alternative splice variants across brain regions and among neuron cell-types (Aoto *et al*., 2013; Fuccillo *et al*., 2015; Földy *et al*., 2016; Lukacsovich *et al*., 2019). These differential expression profiles and their combinatorial interactions with postsynaptic ligands are thought to specify excitatory and inhibitory synapse formation and confer distinct function (Aoto *et al*., 2015; Südhof, 2017; Dai *et al*., 2019). A recent study observed sex-dependent changes in Nrxn3 expression and alternative splicing in mouse hippocampus following chronic stress, suggesting Nrxn3 may engage in sex-specific functions in hippocampus (Freire-Cobo and Wang, 2020). However, previous studies focusing on functional characterization of neurexins have not considered sex as an independent variable.

In hippocampus, Nrxn3 mRNA is highly expressed in GABAergic neurons and exhibits differential isoform expression in distinct interneuron classes (Ullrich *et al*. 1995; Fuccillo *et al.,* 2015). Additionally, neuroligin-2, a prototypical neurexin ligand localized to inhibitory synapses, exhibits the highest binding affinity for Nrxn3β *in vitro* (Koehnke *et al*., 2010). Thus, it was surprising that manipulation of Nrxn3 in developing dissociated hippocampal culture did not alter inhibitory synaptic transmission (Aoto *et al*., 2013; Restrepo *et al*., 2019). Although untested, Nrxn3 may instead impart functional relevance to mature intact hippocampal inhibitory microcircuits where cell-type specific connectivity is preserved. Unique from *NRXN1* and *2*, *NRXN3* mutations are primarily associated with SUDs, SCZ and stress disorders in humans and animal models (Liu *et al*., 2006; Hishimoto *et al*., 2007; Lachman *et al*., 2007; Kelai *et al*., 2008; Novak *et al*., 2009; Brown *et al*., 2011) suggesting that Nrxn3 may dominantly and sex-specifically regulate synaptic properties of neural circuits implicated in these disorders, such as ventral subiculum (vSUB).

vSUB is a understudied subregion of the hippocampal formation that is integral to the regulation of stress, fear, and reward behaviors via its projections to cortical and subcortical regions (Grace, 2010). vSUB hyperexcitability is commonly observed in SCZ, SUDs and depression in animal models and human patients, and is thought to arise from fast-spiking parvalbumin-expressing (PV) interneuron dysfunction (Grace, 2010; Konradi *et al*., 2011; Gill and Grace, 2014). Of interest, PV neurons in hippocampus express Nrxn3 at significantly higher levels than Nrxn-1 and −2 (Nguyen *et al*., 2016; Que *et al*., 2021), suggesting that Nrxn3 may perform a critical, yet unidentified, role in establishing and defining PV interneuron-dependent inhibition within the circuit. Relative to other hippocampal subregions, where local inhibitory architecture is well-defined from decades of studies in male rodents, the inhibitory cell types and patterns of connectivity in vSUB have remained largely enigmatic (Böhm *et al*., 2018). However, the properties and connectivity patterns of subicular regular-spiking (RS) and burst-spiking (BS) pyramidal neurons have been more thoroughly described. RS and BS neurons are discrete classes of pyramidal neurons, which possess striking differences in physiology and brain connectivity, and likely mediate distinct behaviors (Wozny *et al*., 2008; Kim and Spruston, 2011; Graves *et al*., 2012; Cembrowski *et al*., 2018; Wee and MacAskill, 2020). Determining the functional organization of inhibitory input onto RS and BS neurons and investigating what role Nrxn3 plays at vSUB PV inhibitory synapses may elucidate mechanisms that underlie aberrant vSUB activity observed in SCZ and SUDs.

Here we uncover unexpected sexually dimorphic patterning of PV connections in vSUB and identify a novel role for Nrxn3 in mediating inhibitory synaptic transmission in a sex- and cell-type-specific manner. Using optogenetics, paired recordings and imaging, we show that PVs preferentially synapse onto RS neurons in males but BS neurons in females. We find that Nrxn3 KO in PV neurons drastically reduces PV IPSCs in RS neurons in males, driven by a reduction in PV synapse number, but leads to a robust enhancement of IPSCs in females, facilitated by an increase in release probability. Lastly, using single-cell RNA sequencing, we find high expression levels of Nrxn3α and Nrxn1γ in PV neurons, but despite sex-specific Nrxn3 KO effects, detect no sex differences in Nrxn alternative splicing or isoform expression in vSUB PV neurons.

## Results

### The characterization of inhibition in ventral subiculum identifies a bias for burst spiking neurons

The identities and connectivity patterns of vSUB inhibitory neurons that govern principal neuron activity are poorly understood. Thus, to first establish the organization of the inhibitory vSUB microcircuit in its basal state, we asked whether RS and BS neurons are under similar or differential inhibitory control. To address this, we prepared acute *ex-vivo* slices from male and female mice and performed sequential whole-cell recordings from electrophysiologically identified RS and BS neurons (Figure 1A-C). We used electrical stimulation, which non-selectively recruits local inhibitory neurons, and measured the input-output (I/O) relationship and paired-pulse ratios (PPRs) of pharmacologically isolated inhibitory post-synaptic currents (IPSCs). Interestingly, we discovered that the evoked IPSC (eIPSC) amplitudes were significantly greater in BS neurons compared to RS neurons by 82% and 37% in males and females, respectively, without changes in presynaptic release probability (Figure 1D-G), indicating that local vSUB inhibition displays an intriguing postsynaptic cell-type preference.

**Figure 1.**
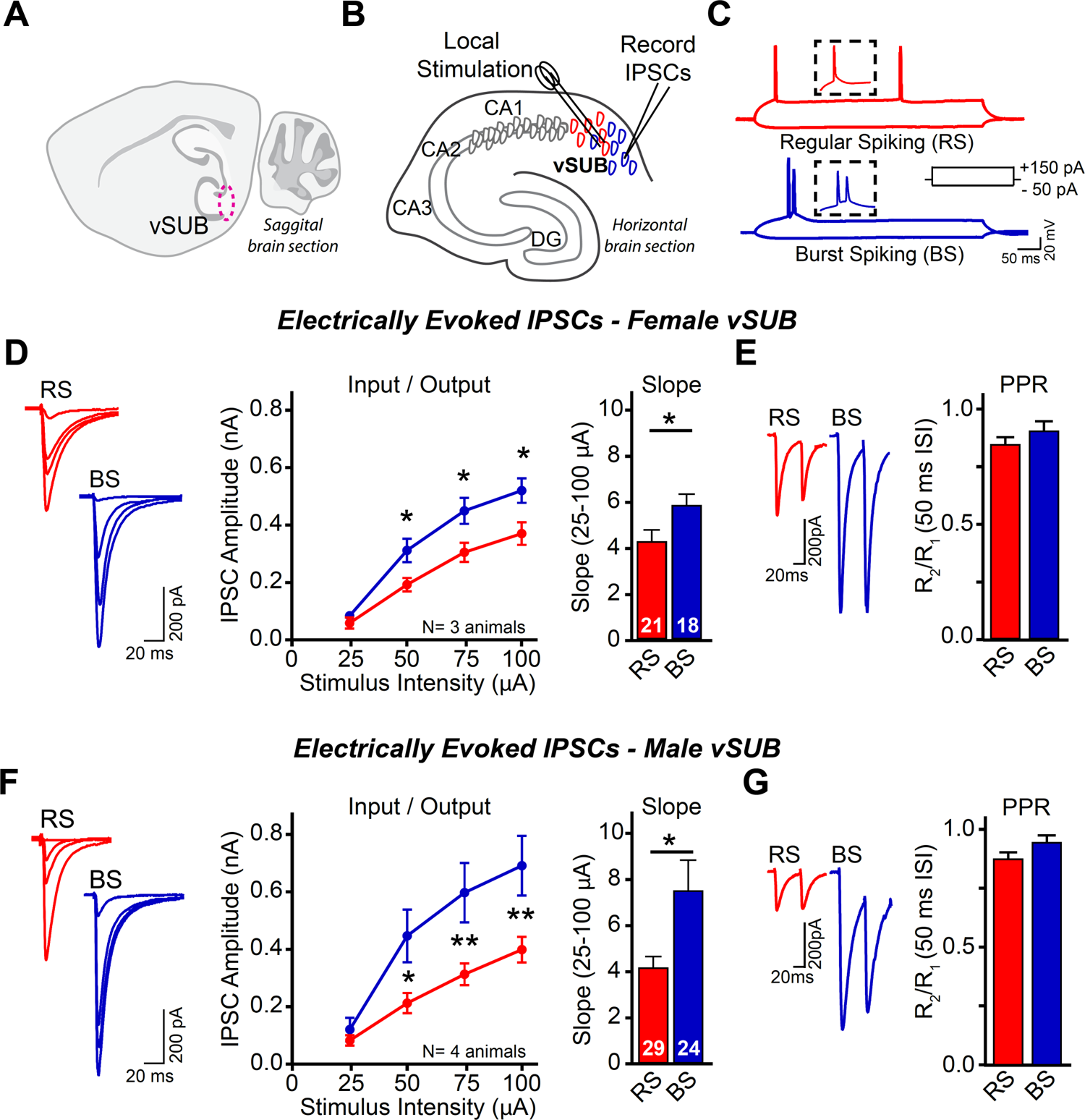
The characterization of inhibition in ventral subiculum identifies a bias for burst spiking neurons. (A) Cartoon of mouse sagittal brain section (circled: ventral subiculum region). (B) Schematic of vSUB local electrical stimulation and whole-cell electrophysiology recordings in horizontal *ex-vivo* hippocampal slices. (C) Identification of regular-(top) and burst-spiking (bottom) cells recorded in current clamp mode. Inset: Expanded view of single vs double (burst) action potentials. (D) Representative traces (left), input-output curves (middle) and summary graph (right) of electrically evoked IPSCs from female RS and BS neurons. n=18-21/3. (E) Representative traces (left) and summary graph (right) of IPSC paired-pulse ratios (50 ms interstimulus interval) measured from RS and BS neurons in female vSUB. (F) Same as (D) but in males. n=24-29/4. (G) Same as (E) but in males. Data points in I/O curves and bar graphs are represented as mean ± SEM; numbers in bars represent number of cells. n=number cells/animals. RS vs BS statistically compared by Student’s t-test at each stimulus intensity in I/O curve, between slopes, and between PPR. *p<0.05; **p<0.01. See also Supplementary Figure 1.

We next asked whether inhibitory postsynaptic strength contributed to the cell-type specific bias in vSUB inhibition. We monitored strontium-evoked asynchronous IPSC (aIPSC) amplitudes in *ex-vivo* slices from RS and BS neurons in male and female mice (Figure S1A-B). aIPSCs are analogous to miniature synaptic events and aIPSC amplitude is thought to reflect postsynaptic strength (Bekkers and Clements, 1999). We found that aIPSC amplitude in BS neurons was ∼25% greater relative to RS neurons in both sexes (Figure S1C-D). Thus, biased inhibition onto BS neurons is at least partially a result of greater postsynaptic strength and not due to differences in presynaptic release probability.

### The usage of Neurexin-3 by inhibitory neurons in ventral subiculum is sexually dimorphic

Next, we asked whether Nrxn3 governs synaptic properties at inhibitory synapses. We injected P21 Neurexin-3α/β^fl/fl^ conditional knock-out mice with active cre-expressing AAV (AAV-hSYN-GFP-Cre; Nrxn3 cKO) into one hemisphere and inactive cre-expressing AAV (AAV-hSYN-mRuby-ΔCre; control) into the other, then performed within-animal assessments by comparing electrically evoked IPSCs from control and cKO slices 14-21 days post-injection (Figure 2A-C). Consistent with our initial findings (Figure 1), eIPSC amplitudes in control slices were larger in BS neurons than RS neurons in both sexes indicating that stereotaxic manipulation does not alter the functional properties of the microcircuit. Following the deletion of Nrxn3, we observed striking sex-specific effects on GABAergic synaptic transmission. In females, genetic ablation of Nrxn3 significantly reduced electrically evoked IPSC amplitudes by 34% in RS and 44% in BS neurons (Figure 2D-E), whereas IPSC amplitudes were unaltered in male Nrxn3 KOs (Figure 2F-G). Independent of sex, deletion of Nrxn3 did not alter IPSC PPRs or IPSC kinetics (Figure S2). Thus, these data unexpectedly reveal that at the same synapses, presynaptic Nrxn3 usage in vSUB inhibitory neurons is sexually dimorphic: in female vSUB, Nrxn3 is required to maintain overall inhibitory strength while in male vSUB, Nrxn3 cKO does not result in a net change in inhibition.

**Figure 2.**
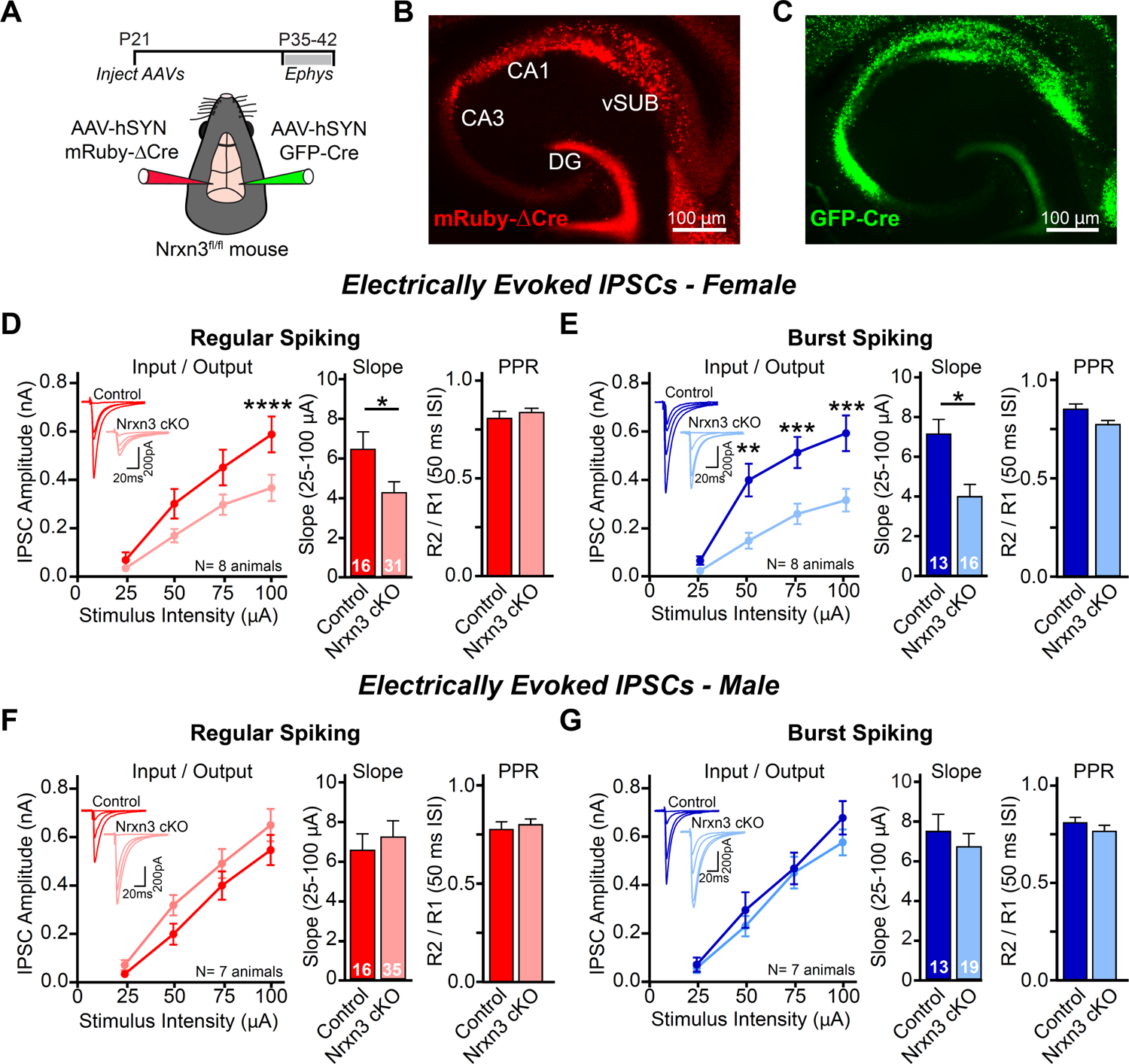
The usage of Neurexin-3 by inhibitory neurons in ventral subiculum is sexually dimorphic. (A) Timeline (top) and schematic of AAV-hSYN-GFP-Cre and AAV-hSYN-mRuby-ΔCre (inactive control) injections into right/left vSUB of Nrxn3α/β conditional KO (cKO) mice, to permit within-animal assessment of Nrxn3 cKO (bottom). (B-C) Representative images of AAV-hSYN-mRuby-ΔCre (left) and AAV-hSYN-GFP-Cre (right) expression in vSUB. (D-E) Ablation of Nrxn3 in female vSUB significantly decreases inhibitory strength at RS and BS synapses. I/O curves and representative traces (inset) (left), summary graphs of I/O slopes (middle) and summary graphs of PPRs (right) of electrically-evoked IPSCs in RS neurons (D) and in BS neurons (E) from control or Nrxn3 cKO slices. (F-G) Same as (D-E) but in males. Unlike females, evoked IPSC amplitudes in RS (F) and BS neurons (G) are unchanged after Nrxn3 cKO. Data points in I/O curves and bar graphs are represented as mean ± SEM; numbers in figures represent number of cells and animals. Control vs cKO statistically compared by Student’s t-test at each stimulus intensity in I/O curve, between slopes, and between PPR. *p<0.05; **p<0.01; ***p<0.001; ****p<0.0001. See also Supplementary Figure 2.

### Parvalbumin interneurons exhibit cell-type and sex-dependent inhibitory bias in the basal subicular microcircuit

While intrigued by the Nrxn3 cKO effects in female vSUB, we were surprised that eIPSCs in males were unaltered given that Nrxn3 is highly expressed in male hippocampal inhibitory neurons (Ullrich *et al.,* 1995; Nguyen *et al*., 2016). However, electrical stimulation evokes neurotransmitter release from mixed GABAergic neuron types. We reasoned that Nrxn3, which displays cell-type specific isoform expression (Fuccillo *et al*., 2015; Lukacsovich *et al*., 2019) may perform divergent and potentially opposing functions in classes of inhibitory interneurons, resulting in no net effect of Nrxn3 cKO on male vSUB IPSCs. To circumvent this potential confound, we interrogated Nrxn3 function in PV interneurons because 1) Nrxn3 is highly expressed in hippocampal PV neurons (Nguyen *et al*., 2016). 2) PVs provide potent inhibitory control of hippocampal principal neuron activity and control sharp-wave ripple oscillatory activity relevant to learning (Schlingloff *et al*., 2014). 3) PV neuron dysfunction contributes to vSUB hyperexcitability observed in SCZ and SUDs - disorders linked to genomic abnormalities in human *NRXN3* (Grace, 2010).

The connectivity and synaptic properties of PV neurons in subicular microcircuitry have received relatively little attention. A study in male mice revealed that PV neurons display a connection bias onto RS cells (Böhm *et al*., 2015), however, synaptic properties of PV neurons in female vSUB is unexplored. To begin to systematically characterize PV inhibition in male and female vSUB, we injected AAV-DIO-ChiEF, a cre-dependent Channelrhodopsin variant, into vSUB of P21 PV-Cre mice and assessed light-evoked PV-IPSCs in *ex-vivo* slices from wild-type RS and BS neurons 14-21 days later (Figure 3A-C). Surprisingly, we found that the functional connectivity of PVs with RS and BS neurons is sexually dimorphic. In female vSUB, the PV IPSC I/O slope was 121% greater for BS neurons compared to RS neurons (Figure 3D). By contrast, in males, PV-IPSC I/O slope was greater for *RS* neurons compared to BS neurons (by 84%), consistent with previous reports (Böhm *et al*., 2015; Figure 3F). Presynaptic release probability was similar at PV-RS and PV-BS synapses in either sex (Figures 3E and 3G). The finding that PV preference for RS neurons in males differs from the BS bias we observed using electrical stimulation (Figure 1) intriguingly suggests that while BS neurons are under stronger *overall* inhibition than RS neurons in both males and females, the interneuron classes that contribute to this inhibitory bias in vSUB differ between sexes.

**Figure 3.**
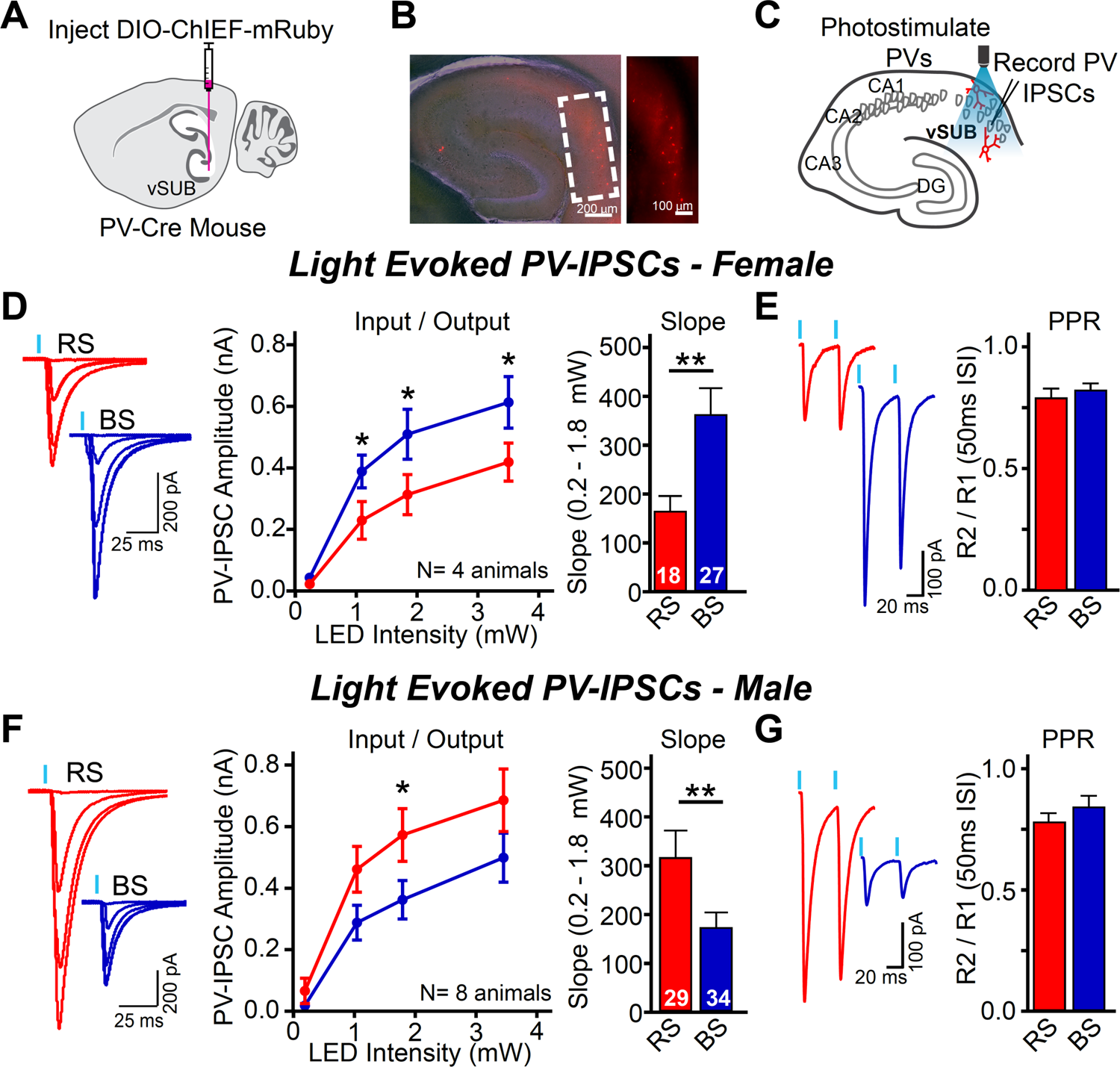
Parvalbumin interneurons exhibit cell-type and sex-dependent inhibitory bias in the basal subicular microcircuit. (A-C) Experimental schematic for isolating PV neuron-mediated synaptic currents. Diagram of AAV injection of Cre-dependent ChIEF into vSUB of PV-Cre mouse (A). Example image of ChiEF-mRuby expression in vSUB PV neurons (overlaid on brightfield image). Inset: magnified vSUB (right) (B). Recording configuration: PV neurons (red) are optogenetically activated with 473nm LED light, eliciting PV-IPSCs recorded in RS/BS neurons (C). (D) In females, PV-synaptic drive is significantly greater onto BS compared to RS neurons. Representative PV-IPSCs (left), input/output (I/O) curves (middle) and corresponding slopes (right) of light-evoked PV-IPSC amplitudes in RS and BS neurons. (E) Representative traces (left) and quantification (right) of light-evoked PV-PPRs from RS and BS neurons. (F-G) Same as D-E but in males. Unlike females, PV-IPSC inhibitory drive is greater onto RS neurons compared to BS neurons. Data points in I/O curves and bar graphs are represented as mean ± SEM; numbers in figures represent number of cells and animals. RS vs BS statistically compared by Student’s t-test at each stimulus intensity in I/O curve, between slopes, and between PPR. *p<0.05; **p<0.01. Note: These data were obtained from control experiments performed in Figure 5. See also Supplementary Figure 3.

### Intrinsic properties of parvalbumin interneurons do not exhibit sexual dimorphism

To test if the sex-specific phenotype was mediated by differences in PV neuron intrinsic excitability, we bred PV-Cre mice to a TdTomato reporter line, Ai9, and then performed whole-cell recordings from labeled PV neurons. We observed no differences in resting membrane properties between females and males (Table S1). We observed a small increase in action potential (AP) half-width in males, but no differences in AP frequency, AP kinetics, rheobase, or sag (Table S1, Figure S3A-D). Moreover, spontaneous excitatory postsynaptic currents were similar between females and males and we found no difference in PV neuron density between male and female vSUB (Figure S3E-F). Therefore, release probability, PV excitability and PV cell number in vSUB are not sexually dimorphic, and likely do not confer the novel sex-specific patterning of PV inhibitory drive among RS/BS neurons. To our knowledge, vSUB has never been described as a sexually dimorphic structure. Intrigued by our unexpected findings, we sought to determine the synaptic properties underlying the basal sex-specific patterning of PV inhibitory drive in detail.

### Sex-specific patterning of vSUB PV-RS and PV-BS preference is mediated by synaptic density in males, quantal size in females

What synaptic properties contribute to the sex dependent and cell-type specific differences in PV-IPSC amplitudes in vSUB? PV-IPSC amplitudes are a product of presynaptic release probabilities, postsynaptic strengths, and number of synaptic contacts. Similar PV-IPSC PPRs in male and female RS and BS neurons indicate release probability is not a factor in the cell-type specific inhibition bias. To examine whether postsynaptic strength and/or synapse numbers are cell-type specific and sexually dimorphic, we utilized strontium-mediated PV-aIPSCs to test for changes in postsynaptic strength and viral labeling to quantify PV synapses.

First, we measured light-evoked strontium-mediated PV-aIPSCs amplitudes in *ex-vivo* slices from male and female PV-Cre mice injected with AAV-DIO-ChIEF (Figure 4A-C). In females, PV-BS aIPSC amplitudes were 30% larger than PV-RS amplitudes (Figure 4D), indicating that PV postsynaptic strength is greater in BS neurons. By contrast, we observed no differences in PV-aIPSC amplitude in males (Figure 4E). To evaluate PV synapse density, we injected a cre-dependent bicistronic AAV expressing a membrane-tethered GFP to label processes and synaptophysin-mRuby to label putative presynaptic sites (AAV-DIO-mGFP-2A-SyPhy-mRuby) (Beier *et al*., 2015) into vSUB of PV-Cre mice (Figure 4F). We filled RS and BS neurons with biocytin, then imaged and quantified the total volume of mRuby+ PV boutons contacting perisomatic regions of 3D reconstructed RS and BS neurons (Figure 4G-H). We quantified total synaptic volume, normalized to the cell fill volume, as a proxy for synapse density, in order to account for overlapping synapses. As previously reported (Harris *et al*., 2001), RS and BS somatic volumes were equal among all conditions (Figure S4). Intriguingly, we found no difference in the density of PV synaptic inputs onto RS and BS neurons in female vSUB (Figure 4I), but quantified more than twice as many PV synapses made onto RS relative to BS neurons in male vSUB (Figure 4J). Together these results reveal that PV interneuron connectivity and basal synaptic properties are sexually dimorphic and cell-type specific: in female vSUB, BS neurons receive greater PV-mediated inhibition due to greater postsynaptic strength, whereas in male vSUB, RS neurons receive greater PV-mediated inhibition due to increased synapse number.

**Figure 4.**
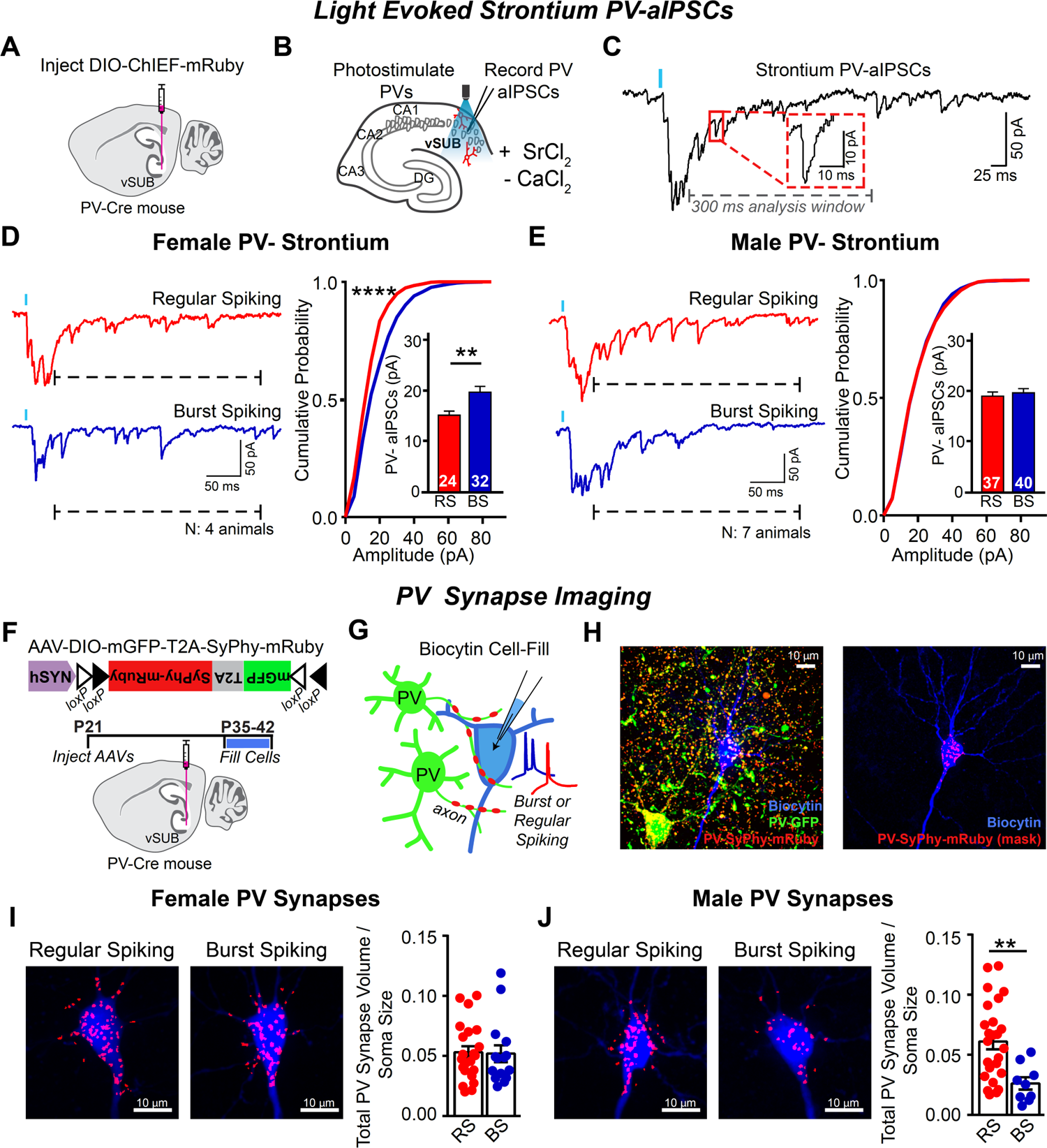
Sex-specific patterning of vSUB PV-RS and PV-BS preference is mediated by synaptic density in males, quantal size in females. (A) Cartoon of AAV injection of Cre-dependent ChIEF into vSUB of PV-Cre mouse. (B) Schematic of recording configuration: Strontium desynchronizes GABA release in PV neurons optogenetically activated with 473nm LED light, eliciting PV-aIPSCs in RS/BS neurons. (C) Example of a light-evoked response in Strontium ACSF. Note asynchronous PV-IPSCs occur for hundreds of milliseconds (analysis window denoted by grey dashed line) following a small phasic PV-IPSC. Red box shows an enlarged, single aIPSC event. (D) PV-aIPSC amplitude is larger in BS compared to RS neurons in females. Representative traces (left), cumulative probability plots (right) of PV-aIPSC amplitude distribution (right) and bar graphs of mean PV-aIPSC amplitudes (inset) from RS (red) and BS (blue) neurons. (E) Same as (D) but in males where PV-aIPSC amplitude is equal between RS and BS neurons. (F) Top: Diagram of the cre-dependent AAV utilized for PV-synaptic bouton labeling. Bottom: experimental timeline and schematic of AAV injection into vSUB. (G) Cartoon of mRuby-labeled PV synapses and cell filling in *ex-vivo* slices. PV neurons (expressing GFP and synaptophysin-mRuby) make synaptic contacts onto an electrophysiologically-identified pyramidal neuron in vSUB. Pyramidal neurons are filled with biocytin for visualization. (H) Left: Max-projected image of a principal neuron (blue) and a neighboring PV neuron (green) in a vSUB slice expressing AAV (see above). Right: same image but showing *somatic* PV+mRuby+ puncta masks, which are utilized in the final quantifications (below). (I) Representative max-projected images (left) and summary graphs representing the average total PV synaptic volume normalized to soma volume (right) of PV synapse density on RS and BS somas in females. RS/BS: n=22/15, N= 4. (J) Same as (I) but in males. PV synapse density is significantly greater onto RS compared to BS somas in males. RS/BS: n=25/9, N= 4. Data in bar graphs are represented as mean ± SEM; n=cells, N=animals; numbers in figures represent number of cells and animals. RS vs BS (bar graphs) statistically compared by Student’s t-test or Mann-Whitney (4I). Cumulative probability plots compared using Kolmogorov-Smirnov (KS) test (stars to left of cumulative plots denote statistical significance in KS test). **p<0.01; ****p<0.0001. Note: These data were obtained from control experiments performed in Figure 7. See also Supplementary Figure 4.

### Neurexin-3 exerts sex-dependent and synapse-specific effects at PV-RS and PV-BS synapses in vSUB

Next we asked if Nrxn3 controls PV-mediated synaptic transmission in vSUB. We crossed PV-Cre mice with Nrxn3α/β^fl/fl^ mice (PV-Nrxn3 KO) (Figure 5A) to selectively delete Nrxn3 from PV-expressing neurons at ∼P9 (Taniguchi *et al*., 2011). We injected AAV-DIO-ChIEF into vSUB of PV-Nrxn3 KO mice and PV-Cre control mice (control mice were separately analyzed in Figures 3-4 for basal circuit characterization), then measured light-evoked PV-IPSCs in *ex-vivo* slices (Figure 5B-D). Interestingly, PV-Nrxn3 KO resulted in profoundly distinct and sexually dimorphic phenotypes between RS and BS neurons. In females, PV-Nrxn3 KO at PV-RS synapses resulted in a two-fold enhancement of PV-IPSC I/O slope and increased presynaptic release probability (Figure 5E-F). At PV-BS female synapses, we observed a minor reduction in the I/O relationship concurrent with a paradoxical 13% decrease in PPR (Figure 5G-H). Conversely in males, loss of Nrxn3 at PV-RS synapses reduced synaptic strength by 55% (Figure 5I), whereas PV-BS IPSCs were unaltered (Figure 5K). Deletion of Nrxn3 from PV neurons did not alter PV-IPSC PPRs in males or PV-IPSC kinetics in either sex (Figures 5J, 5L and S5). These unexpected results reveal that Nrxn3 performs distinct and sex-dependent synaptic functions at PV-RS synapses: Nrxn3 KO produced a PV-IPSC gain-of-function phenotype in females and loss-of-function phenotype in males.

**Figure 5.**
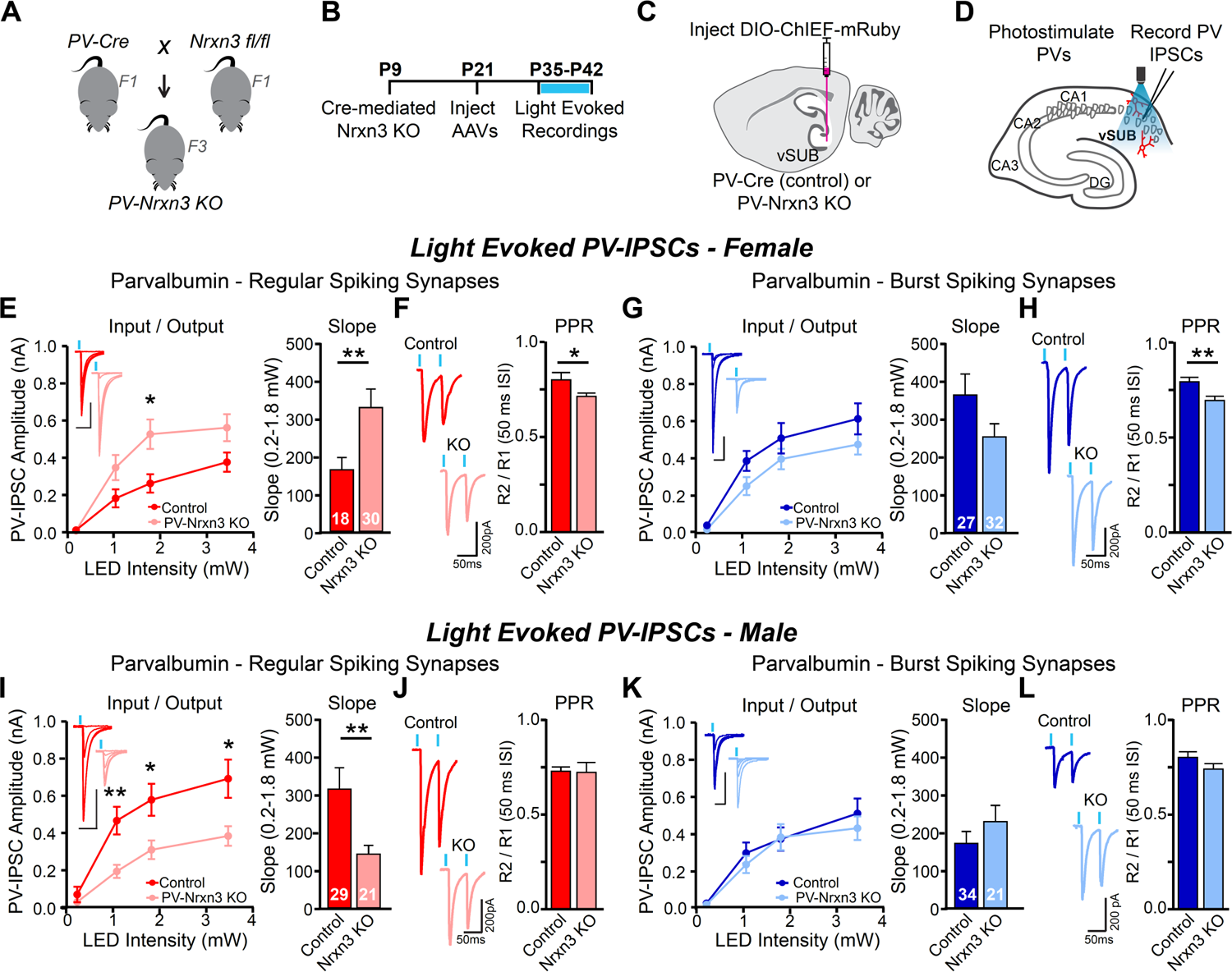
Neurexin-3 exerts sex-dependent and synapse-specific effects at PV-RS and PV-BS synapses in vSUB. (A) Breeding scheme to produce PV-Cre::Nrxn3^fl/fl^ (PV-Nrxn3 KO) mice. (B) Experimental timeline: PV promoter becomes active ∼P9, causing Cre-mediated excision of Nrxn3 in PV-Nrxn3 KO mice. (C) Schematic of AAV injection of DIO ChIEF-mRuby into vSUB of PV-Cre (control) or PV-Nrxn3 KO animals. (D) Schematic of recording configuration: PV neurons are optogenetically activated with 473nm LED light, eliciting PV-IPSCs in RS/BS neurons. (E) PV-Nrxn3 KO enhances female PV-IPSCs in RS neurons. I/O curves (left), representative traces (inset) and slopes (right) of PV-IPSCs in RS neurons from control and PV-Nrxn3 KO animals. (F) Release probability is enhanced in PV-Nrxn3 KO females. Representative traces (left) and quantification (right) of paired-pulse responses at PV-RS synapses in control and PV-Nrxn3 KO animals. (G-H) Same as (E-F) but in BS neurons. PV-IPSCs in BS neurons are not significantly altered in PV-Nrxn3 KO females. (I-J) Same as (E-F) but in males, PV-Nrxn3 KO impairs PV-IPSCs in RS neurons (I) without altering PPR (J). (K-L) Same as (G-H) but in males. PV-Nrxn3 KO does not impact PV-IPSC amplitude (K) or PPR in male BS neurons (L). Data points in I/O and bar graphs are represented as mean ± SEM; numbers in figures represent number of cells. Unlabeled scale bars are 50 ms by 200 pA. N(animals): Female Control= 4; Female KO= 5; Male Control= 8; Male KO= 4. Control vs KO statistically compared by Student’s t-tests. *p<0.05; **p<0.01. Note: the control data obtained from these experiments were also presented in Figure 3 to directly compare RS vs BS for basal circuit characterization. See also Supplementary Figure 5.

### Synaptically-connected paired recordings reveal sex-and synapse-specific roles of PV Nrxn3

To gain greater insight into the role of PV neurons in vSUB microcircuit, we performed paired whole-cell recordings between synaptically coupled presynaptic PV neurons and postsynaptic RS or BS cells in *ex vivo* slices from PV-Cre or PV-Nrxn3 KO. In contrast to our optogenetic approach, paired recordings allow us to precisely monitor unitary PV-IPSCs (uIPSCs) evoked from a single PV interneuron and thus offer a high-resolution assessment of synaptic transmission. We identified presynaptic PV neurons visually and electrophysiologically (Figure 6A-B) then tested the connectivity rate (connected pairs/total pairs tested) and connection strength (uIPSC amplitude, PPR, and failure rate) made onto neighboring principal neurons (Figure 6C). Deletion of Nrxn3 did not alter PV neuron excitability, as intrinsic membrane properties were unchanged (Figure S6A-D).

**Figure 6.**
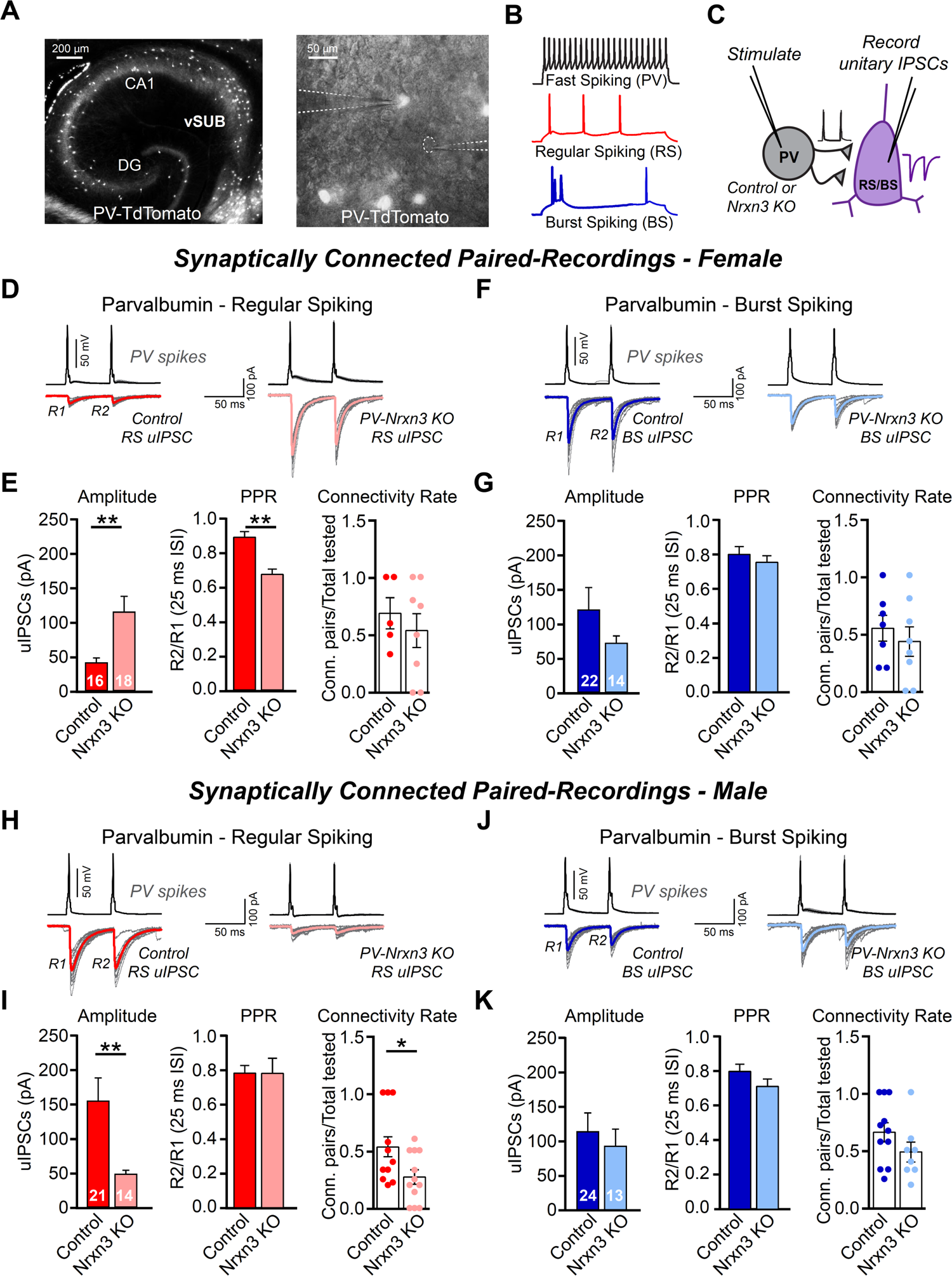
Synaptically-connected paired recordings reveal sex-and synapse-specific roles of PV Nrxn3. (A) Image of TdTomato+ PV neurons in ventral hippocampus from a PV-Cre::Ai9 animal (left) and fluorescent image overlaid on brightfield image of a paired recording (PV neuron-BS neuron) in vSUB of an *ex-vivo* slice (right). (B) Examples of intrinsic firing patterns used to identify fast-spiking PV neurons (top) and regular (middle) and burst spiking (bottom) pyramidal neurons. (C) Illustration of synaptically-connected paired recording: presynaptic PV neurons with intact Nrxn3 (control) or Nrxn3 deletion (PV-Nrxn3 KO) are fired with current injections and resulting PV-unitary IPSCs (uIPSCs) are recorded in postsynaptic RS or BS pyramidal neurons. (D) Representative PPR traces: PV action potentials (black) result in uIPSCs in synaptically-connected RS neurons from PV-Cre (control, left) and PV-Nrxn3 KO female animals (right). Average trace is shown overlaid on multiple trials (grey). (E) uIPSC amplitude and release probability is enhanced in RS neurons in PV-Nrxn3 KO female animals. Quantification of mean uIPSC amplitude (left), PPR (middle), and connectivity rate (right) of PV-RS pairs in control and PV-Nrxn3 KO female animals. (F-G) Same as D-E, but comparing PV-BS pairs from control and PV-Nrxn3 KO female animals. (H-I) Same as (D-E), but in males. PV-Nrxn3 KO results in a loss-of-function phenotype at PV-RS synapses. (J-K) Same as in F-G, but at male PV-BS synapses. Data in bar graphs are represented as mean ± SEM; numbers in bars represent number of pairs; dots on connectivity rate plots represent individual animals. N(animals): Female Control= 8; Female KO= 10; Male Control= 13; Male KO= 15. Control vs KO statistically compared by Student’s t-tests. *p<0.05; **p<0.01. See also Supplementary Figure 6.

Consistent with our optogenetic results, in female PV-RS pairs, PV-Nrxn3 KO resulted in a striking 176% enhancement of uIPSC amplitude, a decrease in PPR and uIPSC failure rate, indicating an increase in presynaptic release probability, and no changes in synaptic connectivity rate (Figures 6D-E and S6E). In female PV-BS pairs, we observed an insignificant decrease in uIPSC amplitudes in PV-Nrxn3 KO animals and no change in either PPR or connectivity rate (Figure 6F-G). Also consistent with our optogenetic data, we observed a significant 69% reduction in uIPSC amplitude at male KO PV-RS synapses (Figure 6H-I). The reduction in uIPSC amplitude was not accompanied by changes in PPR; instead, we found the rate of PV-RS connectivity was significantly reduced (Figure 6I). In male PV-BS pairs, uIPSC amplitudes, PPR and connectivity rate were unaltered (Figure 6J-K). In all conditions, PV synapses exhibited low uIPSC failure rates, indicative of high fidelity of release characteristic of PV interneurons (Bartos and Elgueta, 2012). Ablation of Nrxn3 from PV interneurons did not result in sex-dependent or synapse specific changes to uIPSC kinetics, indicating that Nrxn3 does not control postsynaptic receptor composition at these synapses (Figure S6E-L). Together, the paired recording data reproduced the sexually dimorphic PV-Nrxn3 KO effect on PV-PSC amplitude that we observed using ChIEF stimulation, and excitingly reveal that the gain-of-function PV-IPSC phenotype at female PV-RS synapses is driven by increased release probability whereas the loss-of-function phenotype at male PV-RS synapses might be attributable to changes in synaptic connectivity.

### Nrxn3 controls PV synapse number in a cell-type and sex dependent manner

To further evaluate the synaptic properties controlled by Nrxn3 in PVs, we utilized strontium and imaging experiments detailed previously (Figure 4) to assess synaptic strength and synaptic density in PV-Cre (control) and PV-Nrxn3 KO animals. First, we measured light-evoked, strontium mediated PV-aIPSCs in *ex-vivo* slices. We found that Nrxn3 does not control postsynaptic inhibitory strength onto RS or BS neurons in either sex (Figure 7A-D).

**Figure 7.**
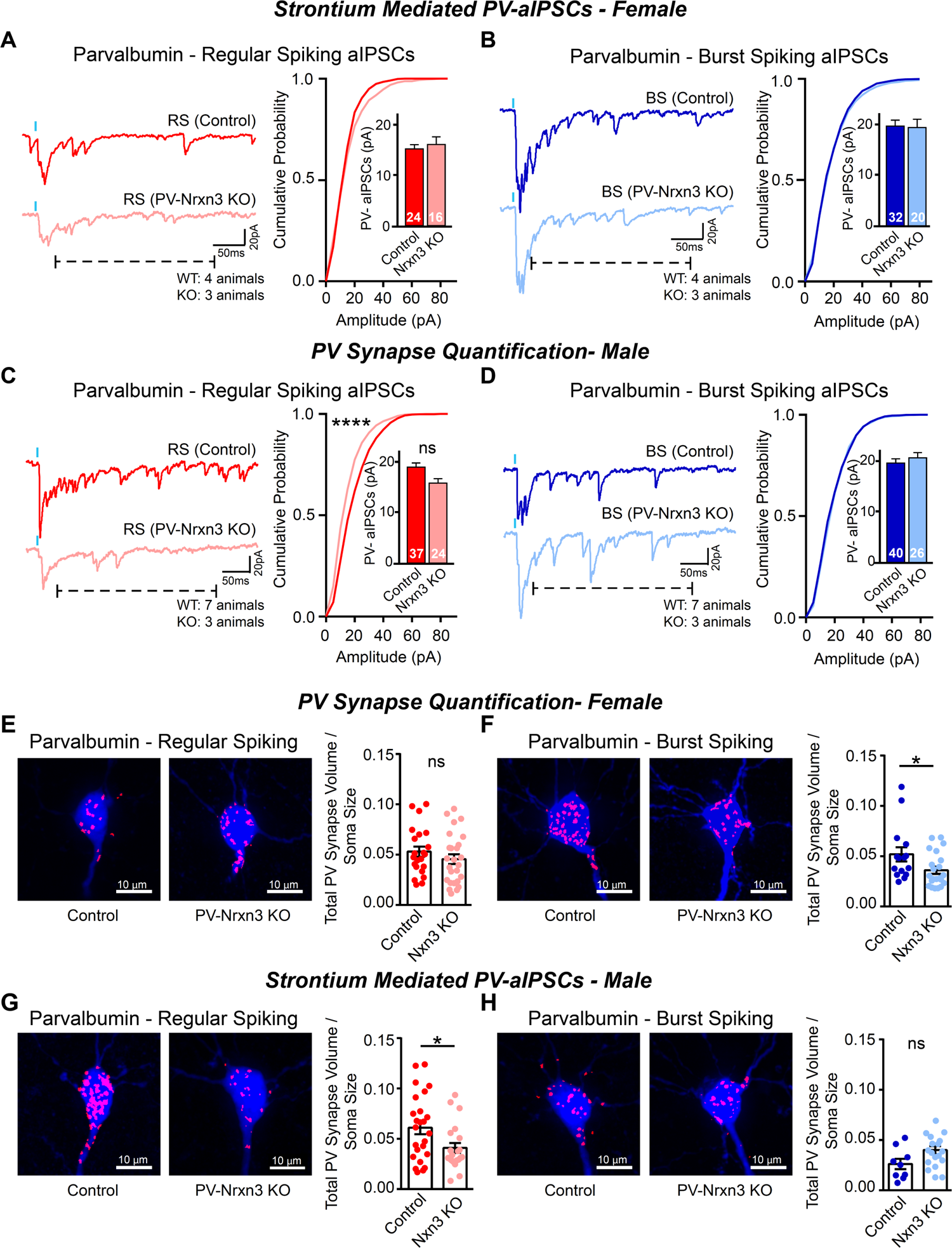
Nrxn3 controls PV synapse number in a cell-type and sex dependent manner. (A-D) Ablation of Nrxn3 from PV neurons does not alter inhibitory postsynaptic strength, measured by light-evoked, strontium-mediated asynchronous IPSCs (PV-aIPSCs), in female (A and B) or male (C and D) RS and BS synapses, respectively. (E-H) PV-Cre (control) and PV-Nrxn3 KO animals were injected with Cre-dependent AAV to label PV synapses (AAV-DIO-mGFP-T2A-Synaptophysin-mRuby) then PV-synapses made onto biocytin filled RS/BS neurons were quantified. Representative, max-projected images (left) and quantification (right) of the average volume of somatic PV-SyPhy-mRuby puncta masks (red) normalized to soma volume of RS and BS cells (blue) from control and PV-Nrxn3 KO females (E-F) and males (G-H). Female Control: RS/BS cells=22/15, N=4 mice; Female KO: RS/BS cells=26/22, N=6 mice. Male Control: RS/BS cells=25/9, N=4 mice; Male KO: RS/BS cells=21/18, N=5 mice. Data in bar graphs are represented as mean ± SEM; numbers in figures represent number of cells and animals. Control vs KO statistically compared using Student’s t-test or Mann-Whitney (7A,C,F). Cumulative probability plots were compared using Kolmogorov-Smirnov (KS) test (stars to left of cumulative plots denote statistical significance in KS test) ns: p>0.05; *p<0.05; ****p<0.0001. Note: the control data from these experiments were also presented in Figure 4 to directly compare RS vs BS for basal circuit characterization. See also Supplementary Figure 4.

Quantification of PV synapses, however, revealed sex- and synapse-specific roles for PV-Nrxn3. We found that Nrxn3 ablation from PV neurons did not alter synapse density on RS neurons in females (Figure 7E), suggesting that increased PV inhibition onto RS neurons is a result of enhanced release probability (Figure 6E). By contrast, at female PV-BS synapses, loss of Nrxn3 produced a significant reduction in synapse density (Figure 7F). Importantly, this effect appears to be partially compensated by the small increase in release probability, which lead to an insignificant reduction in IPSC amplitudes at BS synapses (Figures 5G-H and 6G). In males, consistent with reduced IPSC amplitude and connection probability at PV-RS synapses (Figure 6I), there were significantly fewer PV synapses on RS neurons in KOs compared to control, but no change onto BS neurons (Figure 7G-H). Together, the functional analyses surprisingly revealed Nrxn3 plays pleiotropic roles in PV interneurons in vSUB in a manner dependent on sex and post-synaptic cell-type. At PV-RS synapses, Nrxn3 suppresses PV presynaptic release in females, but promotes synapse density in males. At female PV-BS synapses, similar to male PV-RS synapses, Nrxn3 promotes synapse density but appears to be dispensable at male PV-BS synapses. These findings raise the fascinating possibility that distinct Nrxn3 isoforms or alternative splice variants exhibit sexually dimorphic expression patterns in PV neurons to manifest the disparate functional properties.

### Neurexin isoform expression and alternative splicing in vSUB PVs are comparable between sexes

To investigate Nrxn expression profiles in PV interneurons in male and female vSUB, we harvested fluorescent and electrophysiologically validated single PV neurons from slices made from P35-42 female and male PV-Cre::Ai9 mice and performed single-cell RNA sequencing (Figure 8A). Each cell was inspected for RNA quality, yielded high mapping rates (Figure 8B and S7A) and displayed high levels of parvalbumin expression along with other gene markers typical of PV basket cells (Figure S7C) (Que *et al*., 2021).

**Figure 8.**
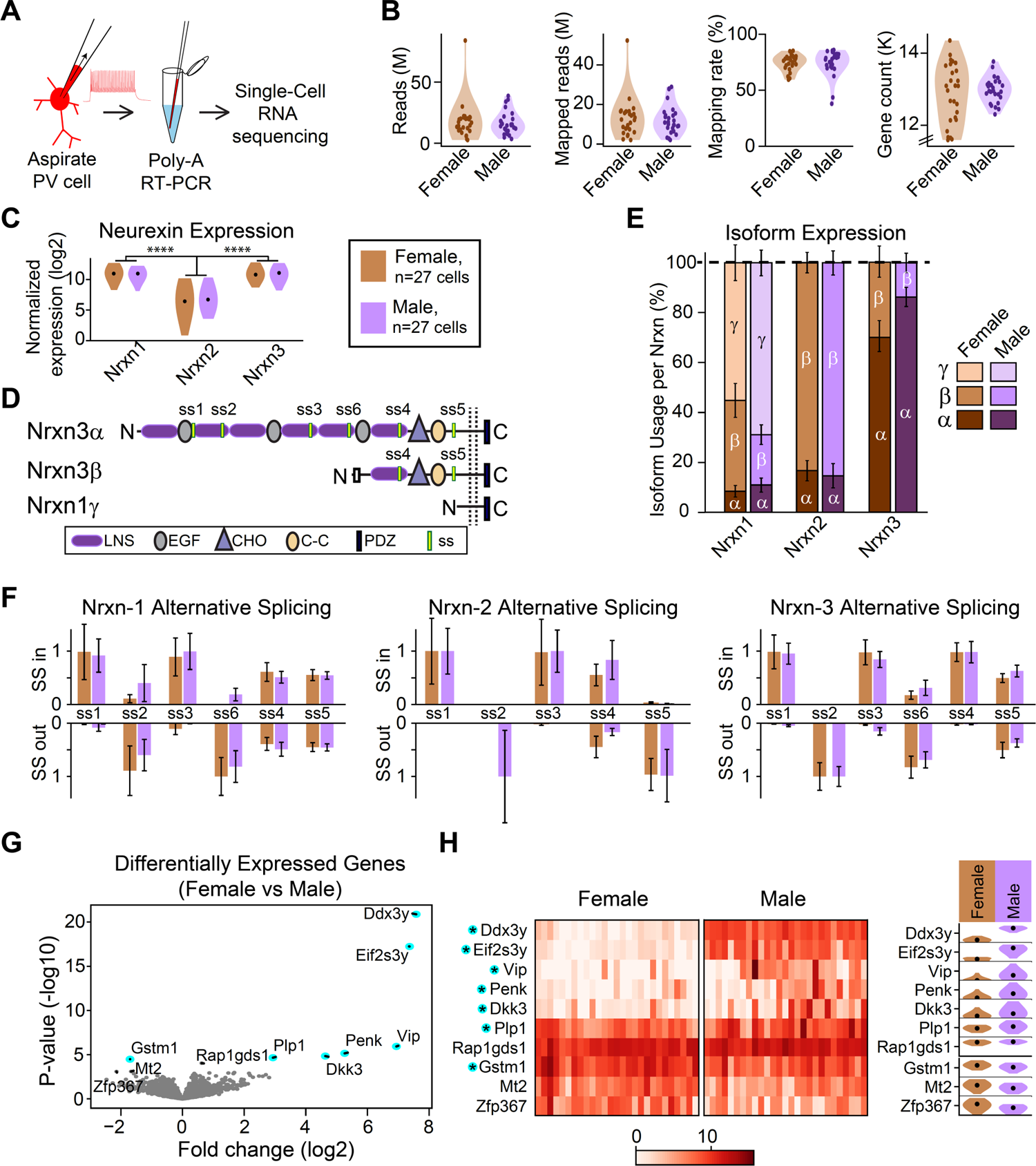
Neurexin isoform expression and alternative splicing in vSUB PVs are comparable between sexes. (A) Experimental steps: TdTomato+ PV neurons were aspirated from male and female vSUB slices then processed individually for RNA sequencing. (B) Violin plots representing the total number of mRNA reads and mapped reads to mouse genome (in millions), mapping rate, and total gene count (in thousands) from female and male PV neurons. (C) Violin plots comparing sce-normalized and log-transformed expression levels of Nrxn1-3 in males and females. Nrxn1 and Nrxn3 are highly expressed compared to Nrxn2, but no differences were found between females vs males. (D) Cartoon of Nrxn3α, Nrxn3β, and Nrxn1-specific γ structures with splice sites (ss) 1-6 labeled. (E) Stacked bar plots showing male/female isoform expression levels as a percentage of total for each Nrxn1-3. Note that while isoform expression varies between Nrxns1-3, relative expression is comparable between females and males. See also Supplementary Fig 7B for sce-normalized isoform expression levels. (F) Bar plots indicating average ratio of splice site (ss) inclusion (up) vs exclusion (down) for Nrxn1 (left), Nrxn2 (middle) and Nrxn3 (right) in males and females. No significant differences were found between male/female. (G) Volcano plot comparing gene expression in female vs male PV neurons with top-ten differentially expressed genes labeled, and statistical significance (FDR<0.05) indicated by blue dot. (H) Heatmap to denote expression levels of each PV cell of the top-ten differentially expressed genes between female vs male (left) and corresponding violin plots (right) showing mean and distribution. The 7 statistically significant genes indicated with blue-highlighted star. Data in bar graphs are represented as mean ± SEM; Female: n=27/3; Male: n=27/4 (cells/mice). Welch tests were used to compare female vs male expression levels for 8C and 8E. In 8C, male and female expression values were combined, then Nrxn 1 vs 2 and Nrxn3 vs 2 expression was compared (Welch test). Quasilikelihood F-test used was used to compare female vs male in 8G. See also Supplementary Figure 7.

Surprisingly, we did not observe significant sex differences in Nrxn expression in vSUB PVs (Figure 8C). Instead, we found high overall levels of Nrxn1 and Nrxn3 and relatively low levels of Nrxn2 expression in vSUB PV cells. High levels of Nrxn1 were unexpected, as other groups have reported Nrxn3 to be the most abundant in CA1 PV cells (Nguyen *et al*., 2016; Que *et al*., 2021). This led us to investigate the expression of Nrxn isoforms in vSUB PV neurons (Figure 8D). We found that the primary Nrxn isoforms expressed are Nrxn1γ and Nrxn3α (Figures 8E and S7B). Nrxn1γ is a highly truncated neurexin isoform with no known role at mammalian synapses (Figure 8D; Sterky *et al*., 2017). By contrast, Nrxn3α contains a full complement of extracellular domains and is able to engage in transsynaptic interactions. Supported by our functional interrogation, this suggests that Nrxn3α dominantly controls synaptic properties of PV neurons in vSUB. We also examined the splice site inclusion rates for Nrxn1-3. We found that alternative exon usage was similar to previous reports from cortex and CA1, indicating that PV neurons exhibit remarkably consistent splicing across brain regions (Figure 8F; Fuccillo *et al*., 2015; Nguyen *et al*., 2016; Lukacsovich *et al*., 2019; Que *et al*., 2021). Surprisingly, transcriptomes of male and female PV neurons were virtually identical with the exception of 7 differentially expressed genes (Figures 8G-H). Taken together, we found that vSUB PV transcriptomes exhibit striking differences in isoform expression among the three Nrxns, but do not display sex-dependent differences in Nrxn expression profiles. This finding indicates that the sex-specific effects of PV-Nrxn3 KO are driven by other elements of the synaptic environment, such as sex-specific expression of post-synaptic adhesion molecules by RS/BS neurons or differences in intracellular signaling.

## Discussion

Despite decades of intense scrutiny, fundamental insight into the distinct and non-overlapping functions of individual neurexins at inhibitory synapses remains elusive. Based on studies from excitatory synapses, there is growing support for the notion that individual neurexins can participate in non-redundant roles at the same synapse as well as distinct roles at different synapses (Aoto *et al*., 2015; Dai *et al*., 2019). A common yet untested assumption is that the usage of neurexins at synapses is independent of sex. Here, we utilized *in vivo* manipulations of Nrxn3 coupled with cell-type- and synapse-specific morphological and electrophysiological analyses and considered sex as a biological variable to ask if Nrxn3 performs a functionally essential role at inhibitory synapses in ventral subiculum. The findings we report here support three unexpected and significantly important conclusions that may cause us to revise how we view the functional organization of local circuits and usage of Nrxns.

First, although local vSUB circuit dysfunction has been implicated in disease, a cell-type-specific understanding of this circuit is limited. Additionally, consistent with most microcircuit dissections in other brain regions, characterization of the vSUB local circuit has only used male animals. Here, we reveal not only that regular- and burst-spiking principal neurons receive different relative levels of overall inhibition, but that the functional organization of PV interneurons in vSUB is sexually dimorphic. In female vSUB, PV interneurons display stronger evoked inhibition onto BS principal neurons. In male vSUB, we observe the opposite - PV interneurons provide greater inhibition onto RS neurons. The most parsimonious explanation of these sex-differences is that they are driven by a common mechanism, however, we discovered that the degree of PV-mediated inhibition is governed by distinct mechanisms. Differences in postsynaptic strength in females and synapse numbers in males drive asymmetric PV inhibition. The sexually dimorphic and cell-type-specific organization of the vSUB microcircuit provides support to the notion that information gleaned from circuit-level studies in male rodents might not always be directly transposable between sexes.

Second, the investigation of Nrxns has historically not considered sex as a biological variable. This has led to the assumption that neurexin function in neurons from male and female rodents are superimposable. Here, we report that conditional deletion of Nrxn3 from PV interneurons resulted in striking sex-dependent synaptic phenotypes at subicular PV-RS synapses. Our data suggest that at female PV-RS synapses, Nrxn3 serves to suppress presynaptic release. By contrast, Nrxn3 promotes synapse maintenance at the same synapses in males. To the best of our knowledge, we provide the first evidence that neurexin usage can be sexually dimorphic.

Third, we report that Nrxn3 expressed in the same presynaptic cell-type can, dependent on postsynaptic cell identity, differentially govern aspects of inhibitory synapse function. In both male and female vSUB, genetic ablation of Nrxn3 in PV positive interneurons resulted in altered inhibition onto RS principal neurons but appeared to be rather dispensable at PV-BS synapses. How can the differential usage of Nrxn3 by PV neurons be explained? Two appealing and non-mutually exclusive possibilities could involve 1) directed localization of Nrxn3α primarily to PV-RS synapses and/or 2) differential localization/expression of inhibitory postsynaptic ligands (e.g. dystroglycan, neuroligin-2 or neuroligin-3) in RS and BS neurons. Our findings suggest that an individual neurexin may not always govern a generalizable function at all synapses made by the same presynaptic neuron and may have broad implications when investigating the circuit-level role of neurexins.

While sex differences in neuromodulation at inhibitory synapses at have been described in hippocampus (Huang and Woolley, 2012; Tabatadze *et al*., 2015), as far as we are aware, our findings represent the first observation of a sex-dependent basal intrinsic connectivity bias. Here, we did not assess variation in circulating gonadal hormones levels among experimental animals. While circulating hormones may play a role in the sex-specific inhibition, it is important to note that the variance in IPSC amplitudes was not significantly different between females and males, suggesting that any hormonal fluctuations among individual animals was not a significant contributing factor mediating the robust phenotypes we observed.

The asymmetric inhibitory control of RS/BS by PVs could facilitate differential engagement of local subiculum and downstream brain regions during specific behaviors. Accordingly, RS and BS neurons are differentially engaged during *in vivo* sharp-wave ripple activity in males, suggesting that inhibitory connectivity bias confers physiological relevance during memory consolidation (Böhm *et al*., 2015; Maslarova *et al*., 2015). The sex differences in PV connectivity and Nrxn3 usage are particularly intriguing in the context of neuropsychiatric disorders. Mutations in *NRXN3* may contribute to sex-specific symptoms by differentially altering vSUB activity in males vs females. Further studies to investigate if vSUB microcircuit connectivity or output is altered sex-specifically in neuropsychiatric disorders would elucidate disease mechanisms and aid therapeutic development. Additionally, it is important to determine whether the sex-dependent organization and usage of Nrxns is a generalizable feature of all microcircuits or specific to a subset of local circuits.

Our single cell RNA-sequencing indicates that these functional sex-differences are likely not a result of differential expression of Nrxn1, 2 or 3 in male vs female PV neurons. Nrxn3’s synaptic roles may be determined by other sex-specific elements of the PV synaptic environment, such as pre-or post-synaptic signaling molecules or usage postsynaptic Nrxn ligands. Importantly, our analyses of neurexin expression relied on quantitative mRNA measurements because antibodies to individual neurexin isoforms are not reliable. To circumvent this limitation, epitope tagged Nrxn1 mice have been generated and have been used to investigate Nrxn1 trafficking (Ribeiro *et al*., 2019) and sub-synaptic localization at excitatory synapses (Trotter *et al*., 2019). However, approaches to investigate the endogenous nanoscale localization of Nrxn3 are undeveloped. Investigating the subsynaptic localization of Nrxn1γ and Nrxn3α at inhibitory synapses could provide critical insights into how these enigmatic molecules contribute to function in vSUB.

## Acknowledgements

We thank members of the Aoto Lab for their thoughtful discussions. This work was supported by grants from the NIH (R00MH103531 and R01 MH116901 to J.A., R35NS116879 to M.J.K. and T32NS099042 to E.E.B.), Brain & Behavior Research Foundation (NARSAD24847 to J.A.), Swiss National Science Foundation (Switzerland, CRETP3_166815 to C.F.).

## Author Contributions

E.E.B. performed animal injections, electrophysiology and synapse imaging, collected and prepared PV mRNA, and performed data analysis. C.S. and D.L. performed RNA sequencing and analysis in the lab of C.F. J.K. performed PV density studies and electrophysiology analysis. S.S. wrote MATLAB code for the PV synapse quantification in the lab of M.J.K. E.E.B. and J.A. are responsible for study conception, experimental design, and data interpretation. E.E.B. and J.A. wrote the manuscript with input from all authors.

## Methods

### RESOURCE AVAILABILITY

#### Lead contact

Further information and requests for resources and reagents should be directed to and will be fulfilled by the lead contact, Jason Aoto.

#### Materials availability

This study did not generate new unique reagents.

#### Data and code availability

The RNA sequencing dataset and MATLAB image quantification code generated during this study are available on NCBI and at Github (https://github.com/samanthalschwartz/Boxer-et-al.git)

### EXPERIEMENTAL MODEL AND SUBJECT DETAILS

Mice were bred at the University of Colorado Anschutz and were from a B6;129 or B6.Cg mixed genetic background. Pvalb^tm1(cre)Arbr^(PV-IRES-Cre) breeders were kindly provided by Dr. Diego Restrepo and *Nrxn3^tm3Sud^*(Nrxn^fl/fl^) breeders were a generous gift from Dr. Thomas Südhof. Gt(ROSA)26Sortm9(CAG-tdTomato)Hze (Ai9) mice and wild-type mice (C57BL/6J) were obtained from The Jackson Laboratory. Mice were housed in a dedicated animal care facility maintained at 35% humidity, 21-23°C, on a 14/10 light/dark cycle. Mice were housed in groups of 2-5 in ventilated cages with same-sex littermates with food and water *ad libitum*. Sex of the animal was determined by external genitalia. Mice were genotyped in-house and all PV-cre+ animals in a littermate were either used for experiments or else randomly chosen to be experimental animals. Animals were stereotactically injected at P21-22, and all other experiments were performed at P35-42 in visibly healthy animals. All procedures were conducted in accordance with guidelines approved by Administrative Panel on Laboratory Animal Care at University of Colorado, Anschutz School of Medicine, accredited by Association for Assessment and Accreditation of Laboratory Animal Care International (AAALAC) (00235).

WT(C57BL/6J)

B6;129-*Nrxn3^tm3Sud^*/J (“Nrxn-3 cKO”: Jax 014157)

B6;129P2-Pvalb^tm1(cre)Arbr^/J homozygote mice (“PV-Cre”: Jax 008069) B6.Cg-Gt(ROSA)26Sortm9(CAG-tdTomato)Hze/J (“Ai9”: Jax 007909)

### METHOD DETAILS

#### Stereotactic Viral Injections

Stereotactic injections were performed on P21-22 mice. Animals were anesthetized with an intraperitoneal injection of 2,2,2-Tribromoethanol (250 mg/kg) then head fixed to a stereotactic frame (KOPF). After drilling small holes in the skull using a handheld drill, 0.5-1.0 µL solutions of adeno associated viruses (AAVs) were injected with pulled glass micropipettes into ventral subiculum at a rate of 11-14 µL/hr using a syringe pump (World Precision Instruments). Coordinates (in mm) were: rostrocaudal: −3.1, mediolateral: +/- 3.2 (relative to Bregma), and dorsoventral: −3.3 (relative to pia). All AAVs used in this study were AAV-DJ vector variants and packaged in-house.

AAV DJ-hSYN-GFP-Cre

AAV DJ-hSYN-mRuby-ΔCre

AAV DJ-hSYN-DIO_loxp_-CHiEF-mRuby

AAV DJ-hSYN-DIO_loxp_-mGFP-T2A-Synaptophysin-mRuby AAV DJ-hSYN-DIO_loxp_-mRuby

#### *Ex-vivo* whole-cell electrophysiology

At P35-P42, animals were deeply anesthetized with isoflurane and decapitated. Brains were rapidly dissected and 300 µm horizontal slices were sectioned with a vibratome (Leica VT1200) in ice cold high-sucrose cutting solution containing (in mM) 85 NaCl, 75 sucrose, 25 D-glucose, 24 NaHCO_3_, 4 MgCl_2_, 2.5 KCl, 1.3 NaH_2_PO_4_, and 0.5 CaCl_2_. Slices were transferred to 31.5°C oxygenated ACSF containing (in mM) 126 NaCl, 26.2 NaHCO_3_, 11 D-Glucose, 2.5 KCl, 2.5 CaCl_2_, 1.3 MgSO_4_-7H_2_O, and 1 NaH_2_PO_4_ for 30 min, then recovered at room temperature for at least 1 hour before recordings. During recordings, slices were superfused with 29.5°C ACSF containing 10 µM NBQX and 50 µM D-AP5 to isolate GABAergic currents, unless otherwise indicated. For strontium experiments, CaCl_2_ was replaced with 2.5mM SrCl_2_ in the ACSF. Pyramidal neurons in subiculum were visually identified with an Olympus BX51W microscope with a 40x dipping objective collected on a Hamamatsu ORCA-Flash 4.0 V3 digital camera using an IR bandpass filter. Neurons were voltage-clamped at −70mV in whole-cell configuration with a high-chloride internal solution containing (in mM) 95 K-gluconate, 50 KCl, 10 HEPES, 10 Phosphocreatine, 4 Mg_2_-ATP, 0.5 Na_2_-GTP, and 0.2 EGTA.

To determine pyramidal neuron identity (regular vs burst spiking), neurons were current-clamped and injected with 500ms depolarizing current in 50 pA steps: cells that fired bursts upon suprathreshold current injection (2-4 spikes with ∼10ms inter-spike interval) were classified as burst spiking whereas those that did not fire bursts were classified as regular spiking. To electrically stimulate local GABAergic cells, a homemade Nichrome bipolar stimulating electrode was gently depressed onto the slice ∼100 µm from the recorded cell and pulsed at 0.1 Hz with a stimulation intensity of 25-100 µA (A-M Systems 2100 Isolated pulse stimulator). To optogenetically stimulate ChiEF expressing PVs, slices were illuminated with 470 nm LED light (ThorLabs M470L2-C1) for 3 ms through the 40x dipping objective located directly over the recorded cell. With an illumination area of 33.18mm^2^ the tissue was excited with an irradiance of 0.006 to 0.17 mW/mm^2^. All recordings were acquired using Molecular Devices Multiclamp 700B amplifier and Digidata 1440 digitizer with Axon pClamp^TM^ 9.0 Clampex software, lowpass filtered at 2 kHz and digitized at 10-20 kHz.

### Synaptically-connected paired recordings

Paired recordings were made from slices prepared in conditions described above; ACSF contained 10 µM NBQX and 50 µM D-AP5 and high chloride internal was used. Td+ PV neurons were patched in whole-cell configuration, current-clamped, and stimulated with 5 ms of 800 pA current to elicit single action potentials. Connectivity rate was calculated as synaptically connected PV-RS or PV-BS pairs divided by the total PV-RS or PV-BS pairs attempted per animal. Animals from which zero connected pairs were achieved were excluded from the calculation, thus actual connection rates are lower than reported.

### Analysis of electrophysiology recordings

Evoked IPSC peak amplitudes from each recording were identified using Axon^TM^ pClamp10 Clampfit software or MATLAB (Mathworks, Inc) using custom scripts. To obtain input/output curves, 12-15 sweeps (0.1 Hz) were averaged to obtain peak amplitude at each stimulus intensity. All final IPSC values are displayed as the absolute value. Input/output slope was calculated using the slope function in excel and was derived from three values representing linear regions of the averaged curve. Release probability was assessed by measurements of paired-pulse ratios (PPRs) at inter-stimulus intervals of 25-55 ms. PPR was measured by dividing the average IPSC amplitude evoked by the second stimulus by the average IPSC amplitude evoked by the first stimulus (R_2_/R_1_). For strontium experiments, slices were evoked at 0.25 Hz 7 times, and amplitudes of asynchronous IPSC events that occurred within a 300 ms window following the phasic IPSC (which occurs due to residual calcium) and amplitude was analyzed using Clampfit event detection software. For analysis of IPSC kinetics, IPSC sweeps were averaged per cell, and rise time and decay tau were calculated using Clampfit functions. Rise time was calculated from the middle 20-80% of the rise slope. Decay time constant values and weights were calculated by fitting the decay slope to a standard, two-term exponential function, fitted with Levenberg-Marquardt method. Weighted Decay Tau was calculated as Tw = ((T1*A1)+(T2*A2))/(A1+A2).

### PV neuron excitability and density

To measure vSUB PV neuron resting membrane and intrinsic firing properties (Supplementary Figure 3), acute slices were made from PV-Cre x Ai9 mice and TdTomato+ cells were patched in whole-cell configuration using an internal solution containing (in mM) 140 K-gluconate, 5 KCl, 10 HEPES, 10 Phosphocreatine, 4 Mg_2_-ATP, 0.5 Na_2_-GTP, and 0.2 EGTA. The ACSF contained 100 µM picrotoxin to isolate spontaneous EPSCs. Cells were first voltage-clamped at −70 to measure EPSCs and resting membrane properties, then current-clamped and given −150 to +400 pA current injections in 25 pA steps. Action potential phase plots, half-width and amplitude were calculated from spikes at rheobase. Sag was calculated as the ratio of the steady state voltage to the maximum decrease in voltage following −150 pA current injection for 250 ms.

To quantify PV cell density in vSUB, PV-Cre x Ai9 mice were transcardially perfused with cold 0.01 M PBS followed by 4% PFA. The brain was fixed in 4% PFA for 2 hours then transferred to a 30% sucrose 4% PFA solution overnight at 4°C. The following day the brain was embedded in OCT compound, rapidly frozen on dry ice, then horizontally cryo-sectioned into 30 µm thick slices. Slices were coverslipped with DAPI fluoromount then imaged with using an Olympus VS120 slide scanning microscope. To quantify PV neuron density, an ROI was drawn around vSUB and TdTomato+ PV cell somas were counted and normalized to ROI area using MATLAB.

### PV synapse imaging

To determine PV synapse density among RS/BS neurons PV-Cre or PV-Nrxn3 KO mice were injected with AAV-hSYN-DIO-mGFP-T2A-Synaptophysin-mRuby into vSUB and 300 µm acute slices were made 14-21 days later. vSUB pyramidal neurons were patched with an internal solution containing biocytin for 15 minutes and fired to determine neuron identity. Slices were then fixed in 4% PFA overnight at 4°C, washed in PBS, then incubated with Cy^TM^5-conjugated Streptavidin (Jackson ImmunoResearch Laboratories, Inc., Lot number 138512) (1:500) in PBS with 0.2% Triton-X for 48 hours at 4°C. Slices were washed and coverslipped then filled cells were imaged on a 3I Marinas spinning disk confocal microscope using a 63x oil objective and z-steps of 0.27-0.5 µm. Images of filled cells were background subtracted, 3D reconstructed and quantified using MATLAB with *DIPimage* toolbox and using custom written routines. Somatic and perisomatic PV synapses were identified as mRuby+, GFP+ puncta that overlapped with Cy-5 signal in the ROI drawn around soma and a ∼10 µm section of apical dendrite. To account for overlapping synapses, total PV synaptic volume colocalized with each RS/BS soma was calculated and normalized to the soma volume as an estimate of synapse density per cell.

### Single-cell mRNA sequencing

Single-cell mRNA was collected from PV neurons and processed using Takara SMART-Seq HT kit according to the manufacturer’s protocol. The work area was wiped down with RNase Away before cell collection. PV cells in vSUB slices were patched in whole-cell configuration using <1 µL of internal solution, fired to confirm fast-spiking identity, and aspirated. Cells were expelled into PCR tubes containing cold lysis buffer and RNase inhibitor, spun briefly, then snap-frozen on dry ice and stored at −80°C until further processing. One-step RTPCR was performed using poly-A selection, eliminating the possibility of genomic contamination. Resulting single-cell cDNA was purified and subsequently analyzed on the Fragment analyzer (Advanced Analytical) to determine concentration and assess sample quality. Library preparation was performed using Nextera XT DNA Library Preparation Kit (Illumina) then sequenced in an Illumina NovaSeq6000 instrument (150 bp, paired-end). Sequencing data was aligned using STAR aligner with the following parameters: trim_front1 = 10, cut_front_mean_quality = 20, average_quality =0, length_required = 30. The quantification was performed with FeatureCounts. Cells were normalized for equal library sizes and differential gene expression analysis was performed using edgeR.

For neurexin isoform level analysis, a modified lasso regression was used to determine the fraction of reads per each exon. Each gene (Nrxn 1-3) was defined by two or three major isoforms: alpha, beta, and gamma, as established in prior literature. An extra factor was allowed for, where each gene had a probability p(ga) of being expressed (depending on the gender and exon), representing alternative splicing, and a penalty factor was added for low p-values. Finally, a decay rate factor was added to account for the potential loss in alignment fidelity farther from the end of an isoform. The predicted isoform expression level then normalized to its single-cell gene expression factor. For alternative splicing analysis, the number of reads that fell on splice junctions were quantified: for each cell, counts of exon inclusion (splice-in) and exon exclusion (splice-out) were added per junction. The mean and SEM for each value, within each sex, was then calculated and normalized such that the two means added to one.

### QUANTIFICATION AND STATISTICAL ANALYSIS

All statistical analysis was performed in Prism 7 (GraphPad) with the exception of strontium cumulative probability plots and single-cell RNA sequencing data. Data were tested for normality using D’Agostino & Pearson normality tests, and Student’s unpaired two-tailed t-tests, Welch tests, or ANOVAs were used to assess differences between groups in normally distributed datasets. If data did not exhibit normal distribution, they were log transformed then analyzed using t-test/AVOVA or assessed using Mann-Whitney or Kruskal-Wallis tests. Male/female Nrxn RNA isoform expression levels were compared using Welch’s t-test. Strontium cumulative probability plots were compared using Kolmogorov-Smirnoff tests on AAT Bioquest (aatbio.com). RNA sequencing analysis was performed using edgeR and Python. Basal sex differences in PV inhibition (Fig 3-4) were determined by analyzing control (wildtype) animals separately from the PV-Nrxn3 KO experiments (Fig 5 & Fig 7) and were presented separately to facilitate clarity and comprehension. All experiments were replicated in at least 3 animals (per condition, per sex). Experimenter was not blinded to animal sex, genotype, or cell identity during data acquisition, but was blinded during image analysis, RNA sequencing analysis, and most electrophysiological analysis. Cells of poor quality were excluded from electrophysiology analysis (i.e. unstable/noisy baseline, access resistance >10% of membrane resistance) and from RNA sequencing (4/58 cells did not pass initial quality control assessments, i.e., cells with library size and expressed gene counts that are three median absolute deviation smaller than the median of library size and median of expressed gene counts from all cells, respectively). Statistical tests used for each experiment are located in figure legends. Sample size can be found in figures and in figure legends and are represented by: N=animal number; n= cell number or pair number. Differences were considered statistically significant if p<0.05. Average values are expressed as mean ± SEM.

## Supplemental Figure Legends

**Supplemental Figure 1 (Relates to Figure 1).**
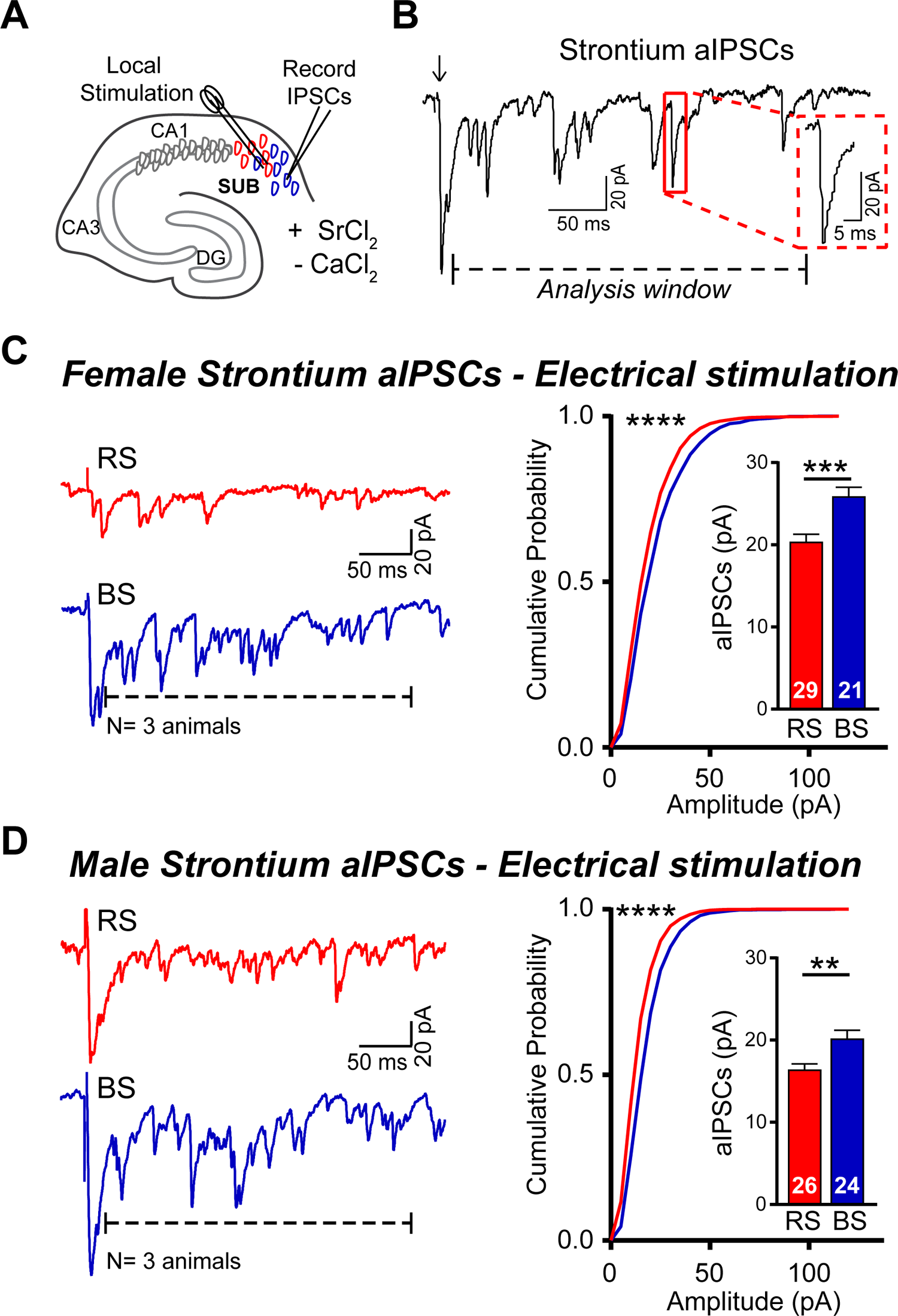
IPSC quantal size is larger in ventral subicular burst spiking neurons compared to regular spiking neurons. (A) Schematic of local electrical stimulation and whole-cell electrophysiology in vSUB in *ex-vivo* hippocampal slices. ACSF calcium is replaced with strontium to produce asynchronous neurotransmitter release. (B) Example of strontium-mediated, electrically-evoked (arrow), asynchronous IPSCs which occur for hundreds of milliseconds following the phasic IPSC. Dashed line shows analysis window (300ms) and a single aIPSC event (red box) is enlarged for detail. (C) Left: Representative strontium-aIPSCs in female RS and BS neurons. Dashed lines denote 300ms analysis windows. Right: Cumulative probability plots showing aIPSC amplitude distribution from female RS (red) and BS (blue) neurons. Inset: summary bar graphs showing larger aIPSC amplitude average in BS neurons compared to RS neurons in females. (D) Same as (C) but in males. Summary data are mean ± SEM; numbers in bars represent number of cells. n=21-29 cells, N=3 mice (female); n=24-26 cells, N=3 mice (male). Student’s t-tests were used to compare BS vs RS (bar graphs). Cumulative probability plots were compared using Kolmogorov-Smirnov (KS) test (stars to left of cumulative plots denote statistical significance in KS test). **p<0.01; ***p<0.001; ****p<0.0001.

**Supplemental Figure 2 (Relates to Figure 2).**
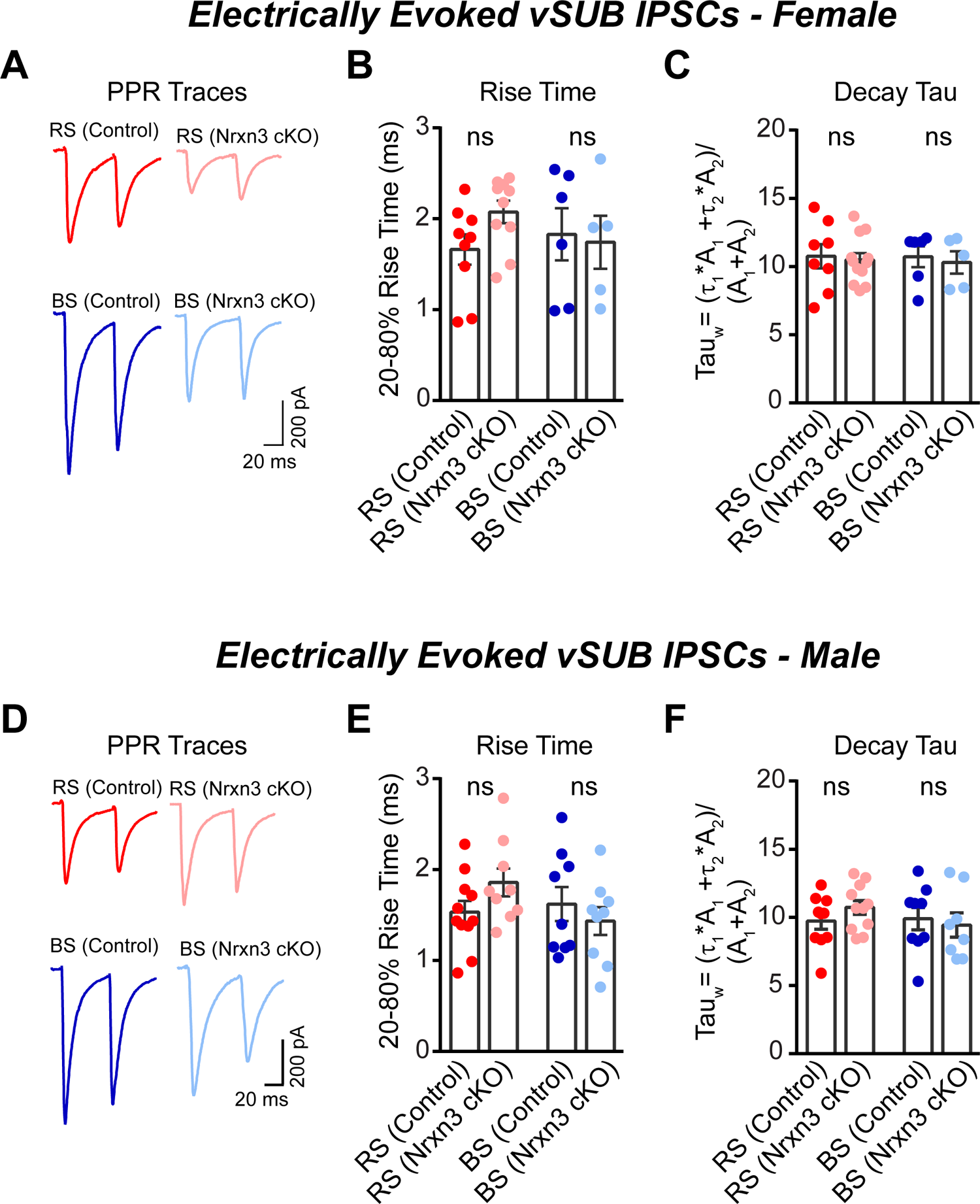
Electrically-evoked IPSC release probability and kinetics in regular and burst spiking neurons are unchanged following Nrxn3 conditional KO in vSUB. (A) Female representative paired-pulse responses to electrical stimulations with 50ms inter-event interval (corresponding data quantified in Figure 2). (B) Female RS and BS IPSC rise time is unchanged following Nrxn3 cKO. (C) Female RS and BS IPSC decay tau (weighted) is unchanged following Nrxn3 cKO. (D-F) Same as (A-C) but in males. Summary data are mean ± SEM; dots on graphs represent individual cells. n=5-8 cells, N=3 mice (female); n=9-11 cells, N=3 mice (male). Student’s t-tests were used to compare control vs cKO. ns: p>0.05.

**Supplemental Figure 3 (Relates to Figure 3).**
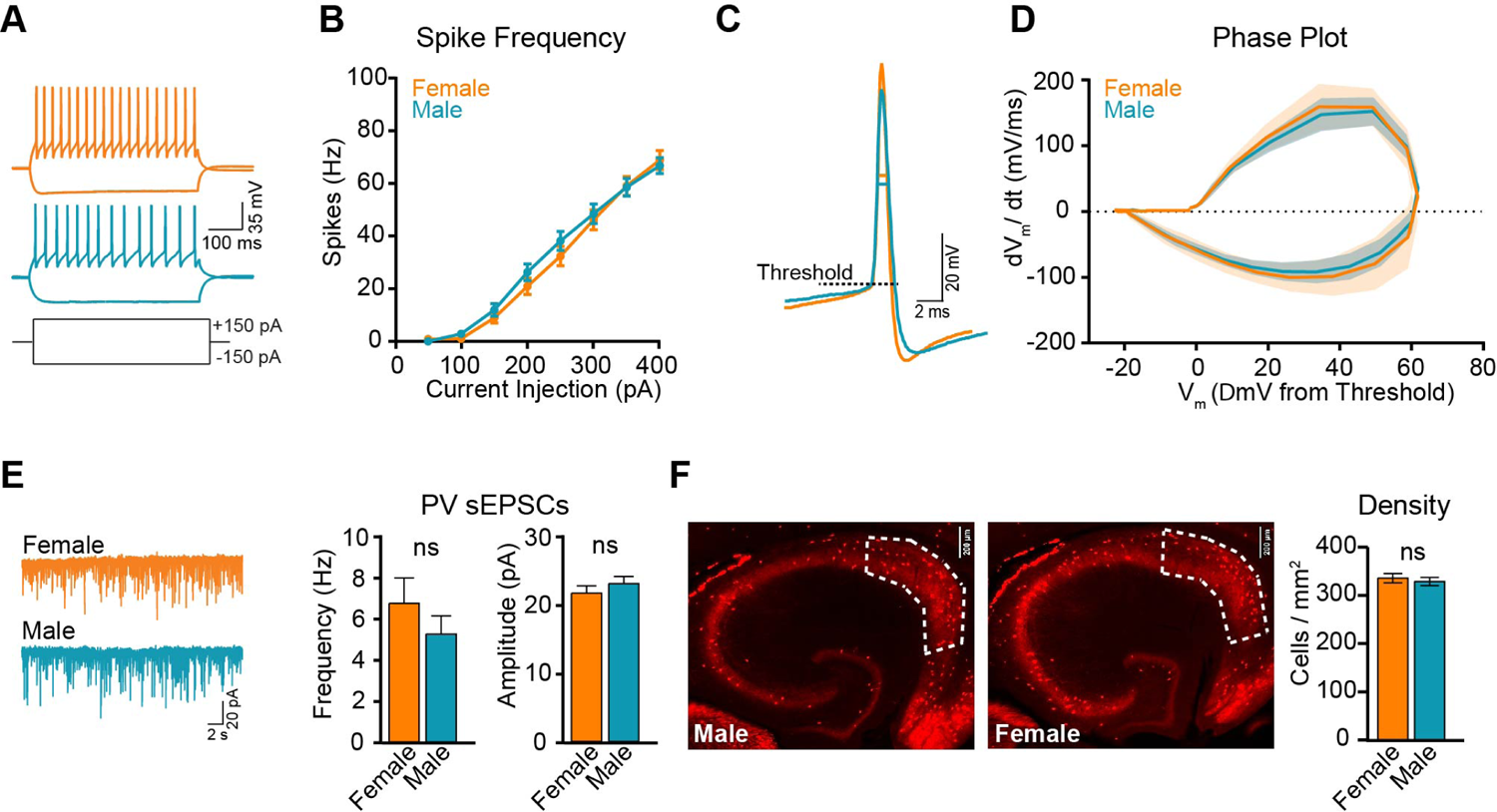
Intrinsic properties of parvalbumin interneurons do not exhibit sexual dimorphism. (A) Examples of physiological responses of female (top) and male (bottom) fast-spiking, Td+ PV interneurons to current injections of 150 and −150 pA. (B) Summary plots of male and female PV neuron spike frequency to increasing levels of current injection. (C) Example traces of single PV action potentials (male and female overlaid) with lines denoting threshold and half-width. (D) Average phase plots produced from single PV action potentials (standard error displayed as lighter color around mean). (E) Representative traces of PV spontaneous EPSCs (left) and bar graphs of average frequency and amplitude (right) of male and female PV sEPSC events. (F) Images of ventral hippocampal slices (30 µm) made from PV-Cre xAi9 mice with vSUB quantification region outlined; Scale bar 200 µm (left).Quantification of Td+ PV neuron density in vSUB region (right). For quantification, averages per animal were calculated, then used to compare male vs female using Student’s t-test. n=78 slices, N=3 mice (female); n=105 slices, N=3 mice (male). Summary data are mean ± SEM. A-E: n=64 cells, N=3 mice (female); n=52 cells, N=3 mice (male). Student’s t-tests were used to compare male vs female. ns: p>0.05. See also Supplementary Table 1.

**Supplemental Figure 4 (Relates to Figures 4 and 7).**
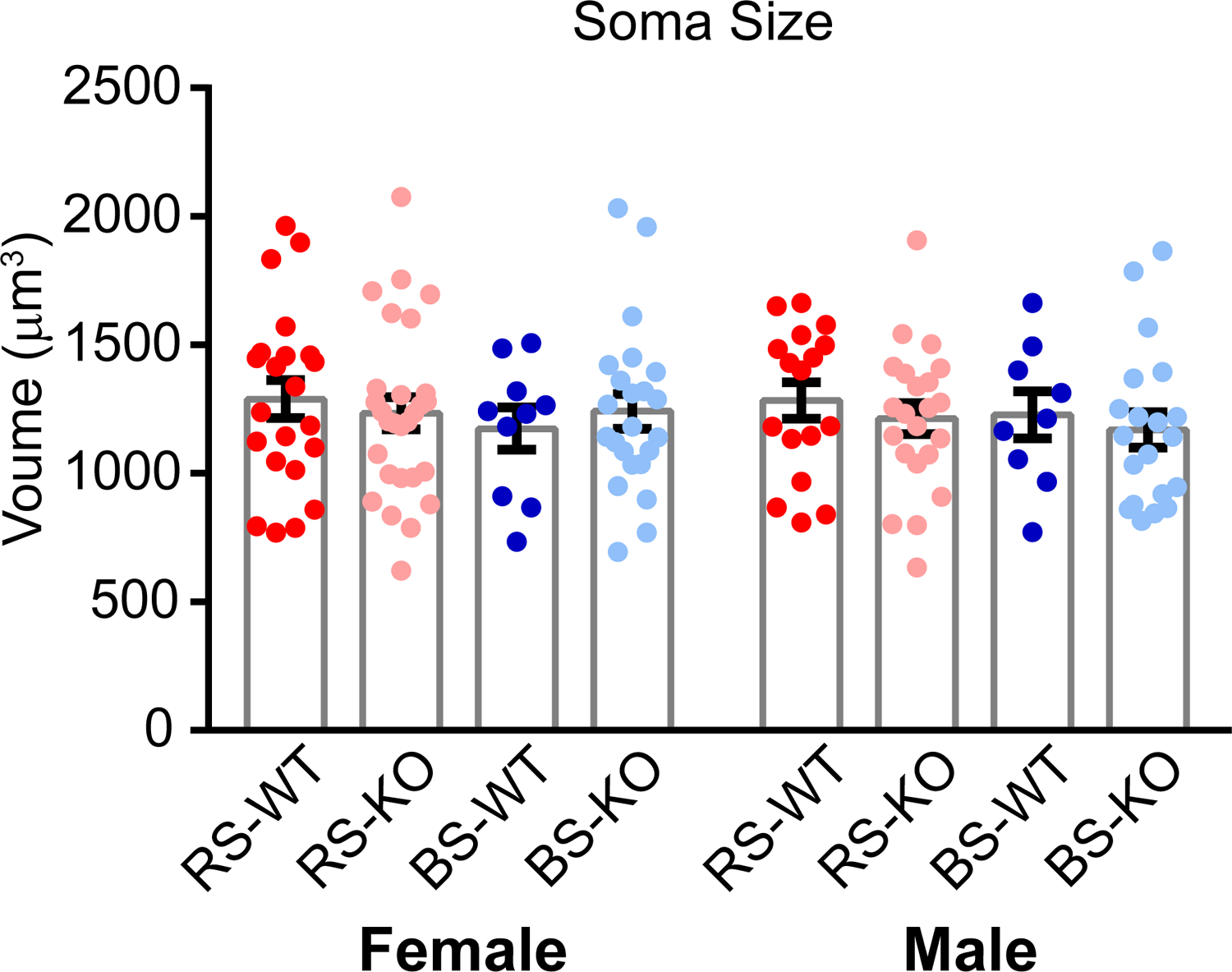
Principal neuron soma size is the same among conditions. (A) Bar charts displaying soma volume means used in PV synapse quantification. Volumes are equal among RS and BS neurons and following PV-Nrxn3 KO in both females and males. Summary data are mean ± SEM. Dots in bars represent cells. n/N=number of cells/mice: Female control: n=22(RS), n=15(BS), N=4; Female KO: n=26(RS), n=22(BS), N=6; Male control: n=25(RS), n=9(BS), N=4; Male KO: n=21(RS), n=18(BS), N=5. ANOVA used to compare female or male means.

**Supplemental Figure 5 (Relates to Figure 5).**
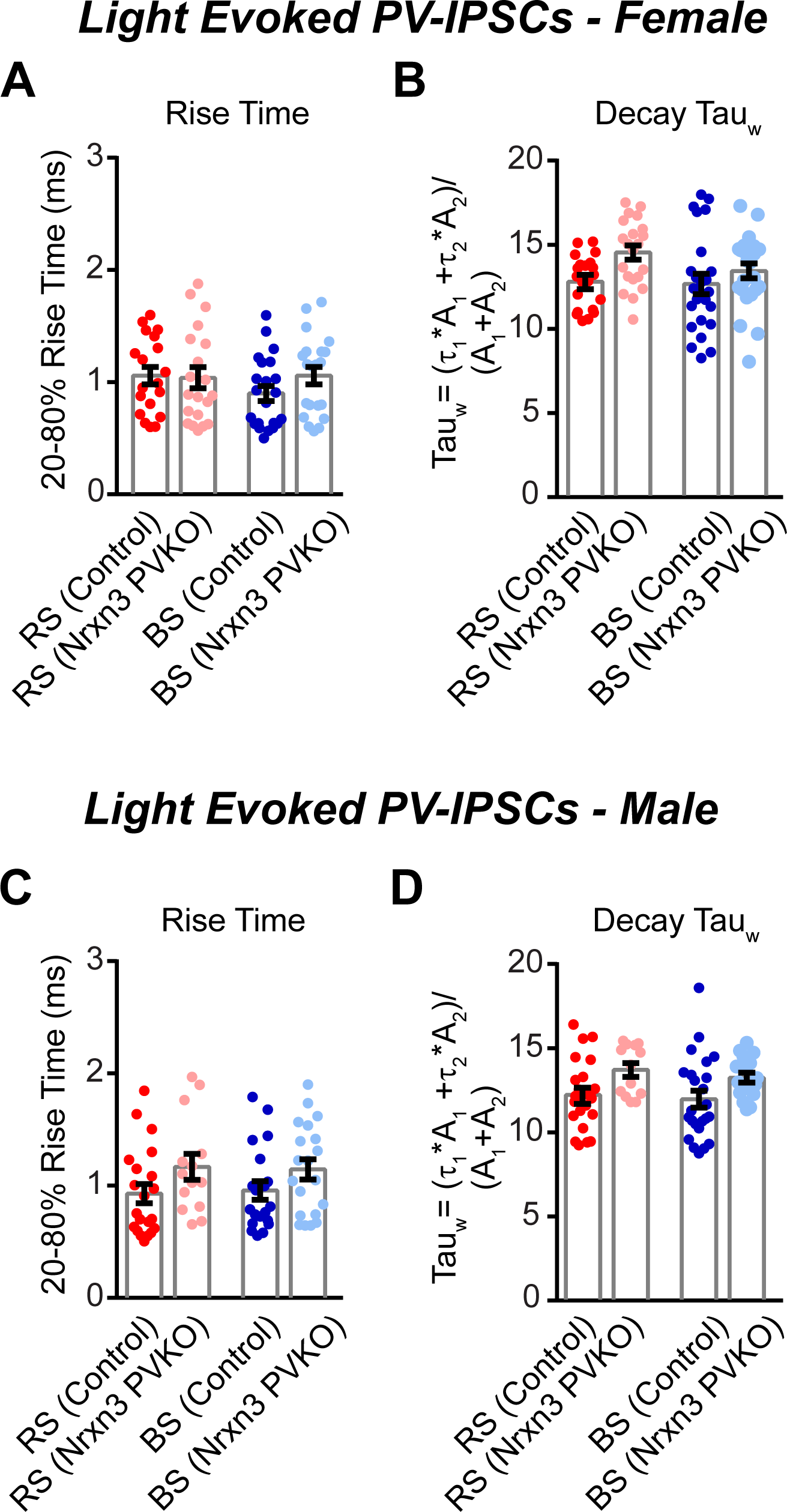
Optically-evoked PV-IPSC kinetics are unchanged following PV-Nrxn3 KO. (A) Female RS and BS PV-IPSC rise time is unchanged following Nrxn3 cKO. (B) Female RS and BS PV-IPSC decay tau (weighted) is unchanged following Nrxn3 cKO. (C-D) Same as (A-B) but in males. Summary data are mean ± SEM. n/N=number of cells/mice: Female control: n=19(RS), n=22(BS), N=5; Female KO: n=20(RS), n=21(BS), N=4. Male control: n=21(RS), n=20(BS), N=4; Male KO: n=14(RS), n=20(BS), N=4. ANOVA used to compare female or male means.

**Supplemental Figure 6 (Relates to Figure 6).**
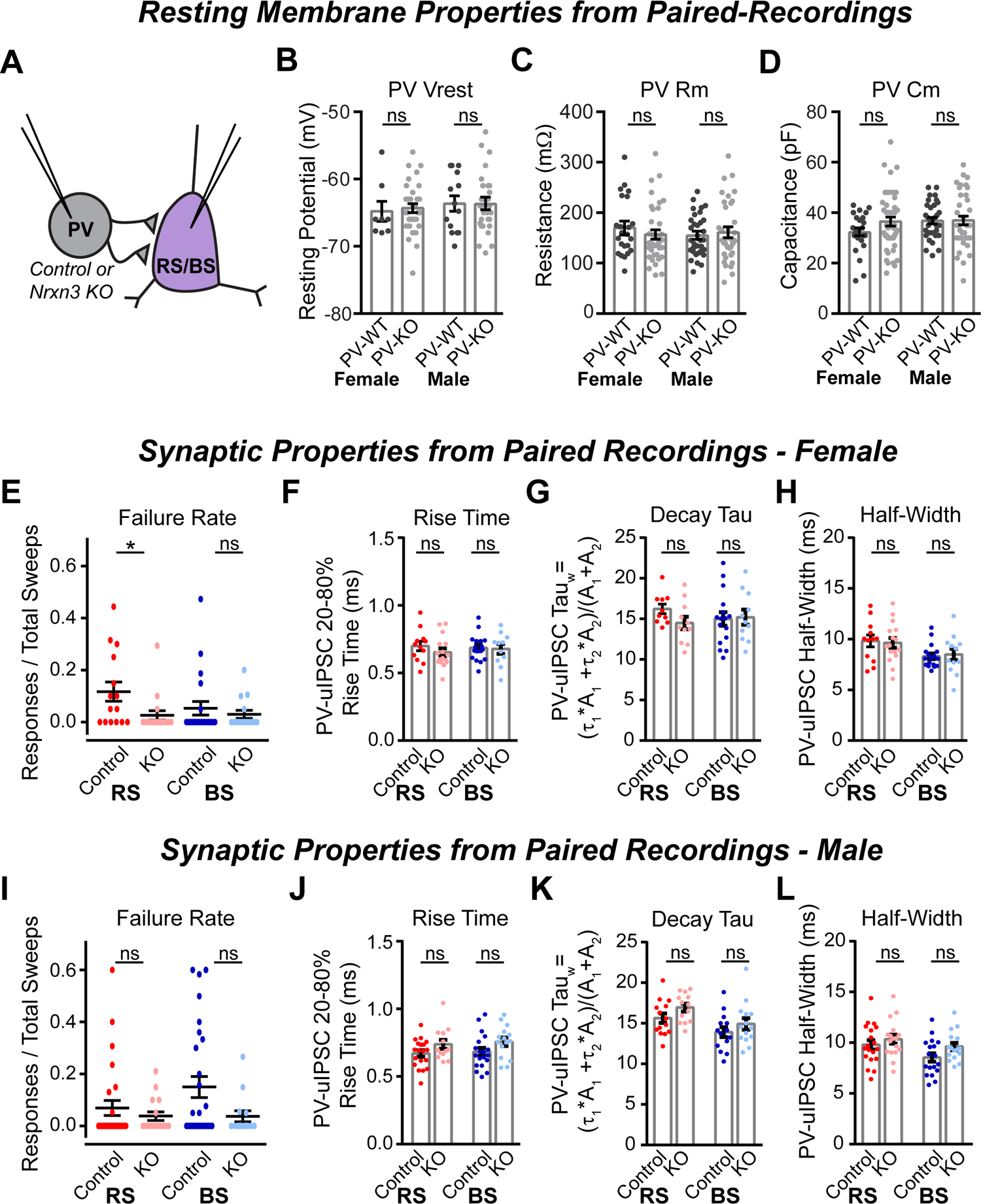
PV interneuron and regular and burst spiking principal neuron intrinsic properties and uIPSC kinetics monitored during paired recordings. (A) Illustration depicting paired recording configuration between presynaptic PV interneuron and postsynaptic regular or burst spiking principal neuron. (B-D) PV intrinsic properties are unaltered following PV-Nrxn3 KO. Bar charts show cell resting potential (B), membrane resistance (C) and capacitance (D) of male and female PV neurons monitored during paired recordings in control and PV-Nrxn3 KO animals. n=23-45 cells. (E) Action-potential-induced GABA release fidelity (failure rate) is reduced in PV-RS but not PV-BS synaptic connections following PV-Nrxn3 KO in females. n=15-21pairs. (F-H) uIPSC kinetics are unaltered following PV-Nrxn3 KO. Bar graphs show rise time (F), weighted decay tau (G) and half-width (H) of uIPSCs in control and PV-Nrxn3 KO females. n=10-17 pairs. (I) Same as (E) but in males. n=14-28 pairs. (J-L) Same as (F-H) but in males. n=10-19 pairs. Graphs show mean; error bars represent SEM. N(animals): Female Control = 8; Female KO = 10; Male Control = 13; Male KO = 15. Control vs KO statistically compared by Student’s t-tests or Mann-Whitney test (S6H, S6L, and S6M-N for BS pairs). ns: p>0.05; *p<0.05

**Supplemental Figure 7 (Relates to Figure 8).**
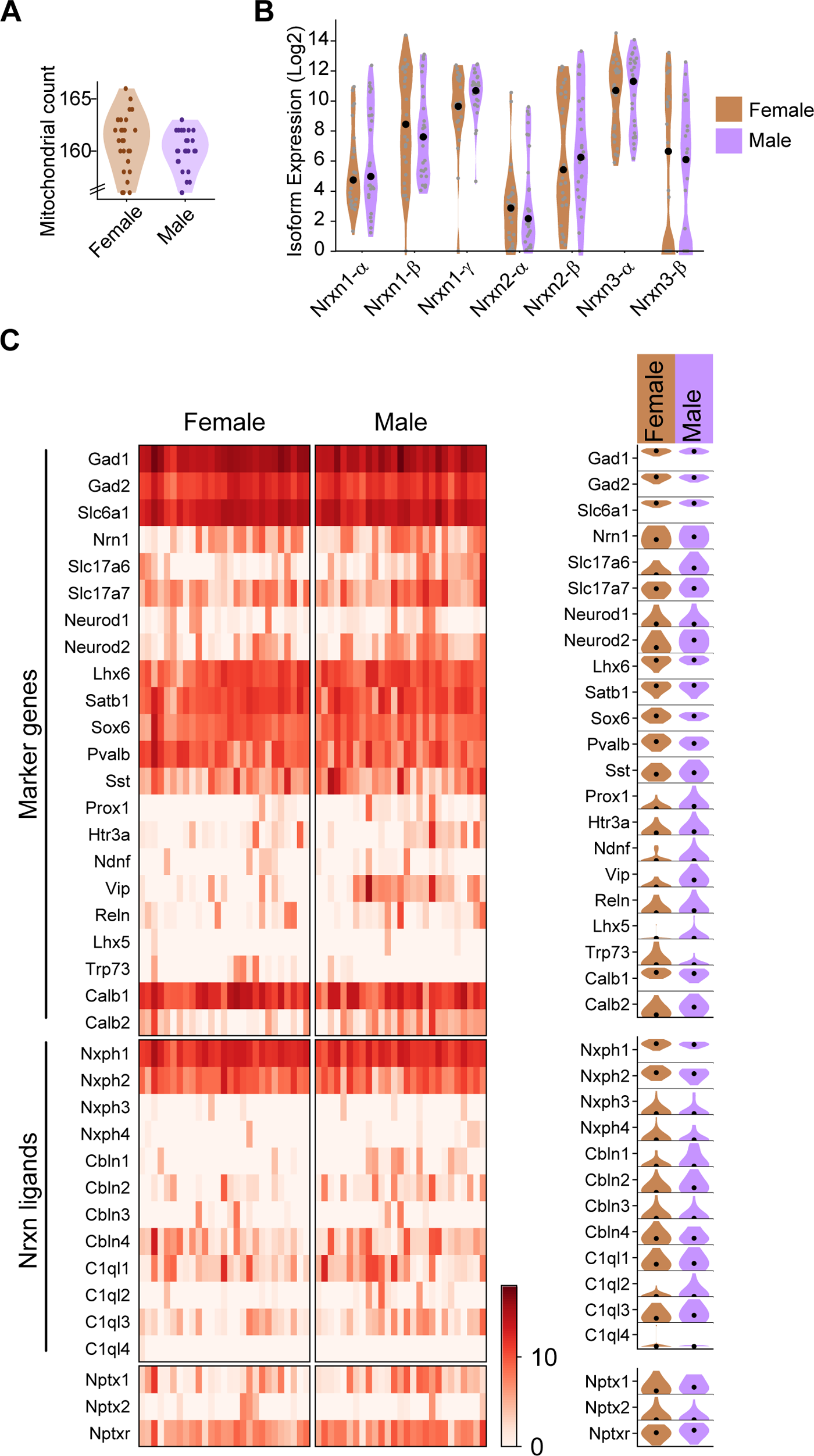
mRNA expression in vSUB PV interneurons determined by single-cell RNA sequencing. (A) Violin plots showing mitochondrial gene count obtained from PV cells. (B) Violin plots showing levels of sce-normalized and log2 transformed Nrxn1-3 isoform expression. (C) Heatmap to denote relative expression levels of each PV cell (left) of canonical PV-marker genes (top) and of Nrxn ligands (bottom) and their corresponding violin plots (right) showing mean and distribution (right) in females and males. Black dots (B, C) represent means. Purple/brown dots (A, B) represent individual cells. n=27 cells, N=3 mice (female); n=27 cells, N=4 mice (male).

**Supplementary Table 1 (Relates to Figure 3 and Supplementary Figure 3).** Intrinsic properties of parvalbumin interneurons measured from whole-cell electrophysiology

|  | **Male**<br>(N= 52 cells,<br>3 animals) | **Female**<br>(N= 64 cells,<br>3 animals) | **Student's**<br>**t-test** |
| --- | --- | --- | --- |
| Resting Potential (mV) | -63.3 ± 0.3 | -63.9 ± 0.2 | p=0.4266 |
| Membrane Resistance (mΩ) | 378 ± 22 | 389 ± 27 | p=0.7545 |
| Capacitance (pF) | 21.1 ± 1.0 | 22.4 ± 1.6 | p=0.4918 |
| AP Half Width (ms) | 0.86 ± 0.02 | 0.77 ± 0.02 | p=0.0010, *** |
| Sag | 1.02 ± 0.02 | 1.02 ± 0.02 | p=0.7350 |
| Rheobase (pA) | 192 ± 10 | 204 ± 9 | p= 0.4130 |

## References

1. Aoto, J., Földy, C., Ilcus, S. M. C., Tabuchi, K., and Südhof, T. C. (2015). Distinct circuit-dependent functions of presynaptic neurexin-3 at GABAergic and glutamatergic synapses. Nature Neuroscience, 18(7), 997–1007. 10.1038/nn.4037

2. Aoto, J., Martinelli, D. C., Malenka, R. C., Tabuchi, K., and Südhof, T. C. (2013). Presynaptic neurexin-3 alternative splicing trans-synaptically controls postsynaptic AMPA receptor trafficking. Cell, 154(1), 75–88. 10.1016/j.cell.2013.05.060

3. Bartos, M., and Elgueta, C. (2012). Functional characteristics of parvalbumin- and cholecystokinin-expressing basket cells. The Journal of Physiology, 590(4), 699–681.10.1113/jphysiol.2011.226175

4. Becker, J. B. (2016). Sex differences in addiction. Dialogues in Clinical Neuroscience, 18(4), 395–402. 10.31887/DCNS.2016.18.4/jbecker.

5. Beier, K. T., Steinberg, E. E., DeLoach, K. E., Xie, S., Miyamichi, K., Schwarz, L., Gao, I. J., Kremer, E. J., Malenka, R. C., Luo, L. (2015). Circuit Architecture of VTA Dopamine Neurons Revealed by Systematic Input-Output Mapping. Cell, 162(3), 622–634. 10.1016/j.cell.2015.07.015

6. Bekkers, J. M., and Clements, J. D. (1999). Quantal amplitude and quantal variance of strontium-induced asynchronous EPSCs in rat dentate granule neurons. The Journal of Physiology, 516(1), 227–248. 10.1111/j.1469-7793.1999.227aa.x

7. Böhm, C., Peng, Y., Geiger, J. R. P., and Schmitz, D. (2018). Routes to, from and within the subiculum. Cell and Tissue Research, 373(3), 557–563. 10.1007/s00441-018-2848-4

8. Böhm, C., Peng, Y., Maier, N., Winterer, J., Poulet, J. F. A., Geiger, J. R. P., and Schmitz, D. (2015). Functional Diversity of Subicular Principal Cells during Hippocampal Ripples. The Journal of Neuroscience, 35(40), 13608–13618. 10.1523/JNEUROSCI.5034-14.2015

9. Boucard, A. A., Chubykin, A. A., Comoletti, D., Taylor, P., and Südhof, T. C. (2005). A Splice Code for Trans-Synaptic Cell Adhesion Mediated by Binding of Neuroligin 1 to alpha- and beta-Neurexins. Neuron, 48(2), 229–236. 10.1016/j.neuron.2005.08.026

10. Brown, S. M., Clapcote, S. J., Millar, J. K., Torrance, H. S., Anderson, S. M., Walker, R., Rampino, A., Roder, J. C., Thomson, P. A., Porteous, D. J., and Evans, K. L. (2011). Synaptic modulators Nrxn1 and Nrxn3 are disregulated in a Disc1 mouse model of schizophrenia. Molecular Psychiatry, 16(6), 585–587. 10.1038/mp.2010.134

11. Cembrowski, M. S., Phillips, M. G., DiLisio, S. F., Shields, B. C., Winnubst, J., Chandrashekar, J., Bas, E., and Spruston, N. (2018). Dissociable Structural and Functional Hippocampal Outputs via Distinct Subiculum Cell Classes. Cell, 173(5), 1280–1292.e18. 10.1016/j.cell.2018.03.031

12. Dai, J., Aoto, J., and Südhof, T. C. (2019). Alternative Splicing of Presynaptic Neurexins Differentially Controls Postsynaptic NMDA and AMPA Receptor Responses. Neuron, 102(5), 993–1008.e5. 10.1016/j.neuron.2019.03.032

13. Földy, C., Darmanis, S., Aoto, J., Malenka, R. C., Quake, S. R., and Südhof, T. C. (2016). Single-cell RNAseq reveals cell adhesion molecule profiles in electrophysiologically defined neurons. Proceedings of the National Academy of Sciences, 113(35), E5222 LP-E5231. 10.1073/pnas.1610155113

14. Freire-Cobo, C., and Wang, J. (2020). Dietary phytochemicals modulate experience-dependent changes in Neurexin gene expression and alternative splicing in mice after chronic variable stress exposure. European Journal of Pharmacology, 883, 173362. 10.1016/j.ejphar.2020.173362

15. Fuccillo, M. V., Földy, C., Gökce, Ö., Rothwell, P. E., Sun, G. L., Malenka, R. C., and Südhof, T. C. (2015). Single-Cell mRNA Profiling Reveals Cell-Type-Specific Expression of Neurexin Isoforms. Neuron, 87(2), 326–340. 10.1016/j.neuron.2015.06.028

16. Gill, K. M., and Grace, A. A. (2014). Corresponding decrease in neuronal markers signals progressive parvalbumin neuron loss in MAM schizophrenia model. The International Journal of Neuropsychopharmacology, 17(10), 1609–1619. 10.1017/S146114571400056X

17. Grace, A. A. (2010). Dopamine system dysregulation by the ventral subiculum as the common pathophysiological basis for schizophrenia psychosis, psychostimulant abuse, and stress. Neurotoxicity Research, 18(3–4), 367–376. 10.1007/s12640-010-9154-6

18. Graves, A. R., Moore, S. J., Bloss, E. B., Mensh, B. D., Kath, W. L., and Spruston, N. (2012). Hippocampal Pyramidal Neurons Comprise Two Distinct Cell Types that Are Countermodulated by Metabotropic Receptors. Neuron, 76(4), 776–789. 10.1016/j.neuron.2012.09.036

19. Harris, E., Witter, M. P., Weinstein, G., and Stewart, M. (2001). Intrinsic connectivity of the rat subiculum: I. Dendritic morphology and patterns of axonal arborization by pyramidal neurons. The Journal of Comparative Neurology, 435(4), 490–505. 10.1002/cne.1046

20. Hishimoto, A., Liu, Q.-R., Drgon, T., Pletnikova, O., Walther, D., Zhu, X.-G., Troncoso, J. C., and Uhl, G. R. (2007). Neurexin 3 polymorphisms are associated with alcohol dependence and altered expression of specific isoforms. Human Molecular Genetics, 16(23), 2880–2891. 10.1093/hmg/ddm247

21. Huang, G. Z., & Woolley, C. S. (2012). Estradiol acutely suppresses inhibition in the hippocampus through a sex-specific endocannabinoid and mGluR dependent mechanism. Neuron, 74(5), 801–808. 10.1016/j.neuron.2012.03.035.

22. Kelai, S., Maussion, G., Noble, F., Boni, C., Ramoz, N., Moalic, J., Peuchmaur, M., Gorwood, P., and Simonneau, M. (2008). Nrxn3 upregulation in the globus pallidus of mice developing cocaine addiction. Neuroreport, 19(7), 3–7. 0.1097/WNR.0b013e3282fda231.

23. Kim, Y., and Spruston, N. (2011). Target-specific output patterns are predicted by the distribution of regular-spiking and bursting pyramidal neurons in the subiculum. Hippocampus, 22(4), 693–706. 10.1002/hipo.20931

24. Koehnke, J., Katsamba, P. S., Ahlsen, G., Bahna, F., Vendome, J., Honig, B., Shapiro, L., and Jin, X. (2010). Splice form dependence of beta-neurexin/neuroligin binding interactions. Neuron, 67(1), 61–74. 10.1016/j.neuron.2010.06.001

25. Konradi, C., Yang, C. K., Zimmerman, E. I., Lohmann, K. M., Gresch, P., Pantazopoulos, H., Berreta, S., and Heckers, S. (2011). Hippocampal interneurons are abnormal in schizophrenia. Schizophrenia Research, 131(1), 165–173. 10.1016/j.schres.2011.06.007

26. Lachman, H. M., Fann, C. S. J., Bartzis, M., Evgrafov, O. V, Rosenthal, R. N., Nunes, E. V, Miner, C., Santana, M., Gaffney, J., Riddick, A., Hsu, C.-L., and Knowles, J. A. (2007). Genomewide suggestive linkage of opioid dependence to chromosome 14q. Human Molecular Genetics, 16(11), 1327–1334. 10.1093/hmg/ddm081

27. Leung M.D., D. A., and Chue M. R. C. Psych., D. P. (2003). Sex differences in schizophrenia, a review of the literature. Acta Psychiatrica Scandinavica, 101(401), 3–38. 10.1111/j.0065-1591.2000.0ap25.x

28. Liu, Q.-R., Drgon, T., Johnson, C., Walther, D., Hess, J., and Uhl, G. R. (2006). Addiction molecular genetics: 639,401 SNP whole genome association identifies many “cell adhesion” genes. American Journal of Medical Genetics Part B: Neuropsychiatric Genetics, 141B(8), 918–925. 10.1002/ajmg.b.30436

29. Lukacsovich, D., Winterer, J., Que, L., Luo, W., Lukacsovich, T., and Földy, C. (2019). Single-Cell RNA-Seq Reveals Developmental Origins and Ontogenetic Stability of Neurexin Alternative Splicing Profiles. Cell Reports, 27(13), 3752–3759.e4. 10.1016/j.celrep.2019.05.090

30. Maslarova, A., Lippmann, K., Salar, S., Rösler, A., and Heinemann, U. (2015). Differential participation of pyramidal cells in generation of spontaneous sharp wave-ripples in the mouse subiculum in vitro. Neurobiology of Learning and Memory, 125, 113–119. 10.1016/j.nlm.2015.08.008

31. Nguyen, T.-M., Schreiner, D., Xiao, L., Traunmüller, L., Bornmann, C., and Scheiffele, P. (2016). An alternative splicing switch shapes neurexin repertoires in principal neurons versus interneurons in the mouse hippocampus. ELife, 5, e22757. 10.7554/eLife.22757

32. Novak, G., Boukhadra, J., Shaikh, S. A., Kennedy, J. L., and Le Foll, B. (2009). Association of a polymorphism in the NRXN3 gene with the degree of smoking in schizophrenia: A preliminary study. The World Journal of Biological Psychiatry, 10(4–3), 929–935. 10.1080/15622970903079499

33. Que, L., Lukacsovich, D., Luo, W., and Földy, C. (2021). Transcriptional and morphological profiling of parvalbumin interneuron subpopulations in the mouse hippocampus. Nature Communications, 12(1), 108. 10.1038/s41467-020-20328-4

34. Restrepo, S., Langer, N. J., Nelson, K. A., and Aoto, J. (2019). Modeling a Neurexin-3α Human Mutation in Mouse Neurons Identifies a Novel Role in the Regulation of Transsynaptic Signaling and Neurotransmitter Release at Excitatory Synapses. The Journal of Neuroscience, 39(46), 9065 LP-9082. 10.1523/JNEUROSCI.1261-19.2019

35. Ribeiro, L. F., Verpoort, B., Nys, J., Vennekens, K. M., Wierda, K. D., and de Wit, J. (2019). SorCS1-mediated sorting in dendrites maintains neurexin axonal surface polarization required for synaptic function. PLoS Biology, 17(10). 10.1371/journal.pbio.3000466

36. Schlingloff, D., Káli, S., Freund, T. F., Hájos, N., and Gulyás, A. I. (2014). Mechanisms of Sharp Wave Initiation and Ripple Generation. The Journal of Neuroscience, 34(34), 11385 LP-11398. 10.1523/JNEUROSCI.0867-14.2014

37. Sterky, F. H., Trotter, J. H., Lee, S.-J., Recktenwald, C. V, Du, X., Zhou, B., Zhou, P., Schwenk, J., Fakler, B., and Südhof, T. C. (2017). Carbonic anhydrase-related protein CA10 is an evolutionarily conserved pan-neurexin ligand. Proceedings of the National Academy of Sciences of the United States of America, 114(7), E1253– E1262. 10.1073/pnas.1621321114

38. Südhof, T. C. (2017). Synaptic Neurexin Complexes: A Molecular Code for the Logic of Neural Circuits. Cell, 171(4), 745–769. 10.1016/j.cell.2017.10.024

39. Tabatadze, N., Huang, G., May, R. M., Jain, A., & Woolley, C. S. (2015). Sex Differences in Molecular Signaling at Inhibitory Synapses in the Hippocampus. The Journal of Neuroscience, 35(32), 11252 LP-11265. 10.1523/JNEUROSCI.1067-15.2015

40. Taniguchi, H., He, M., Wu, P., Kim, S., Paik, R., Sugino, K., Kvitsiani, D., Fu, Y., Lin, Y., et. al., and Huang, Z. J. (2011). A resource of Cre driver lines for genetic targeting of GABAergic neurons in cerebral cortex. Neuron, 71(6), 995–1013. 10.1016/j.neuron.2011.07.026

41. Trotter, J. H., Hao, J., Maxeiner, S., Tsetsenis, T., Liu, Z., Zhuang, X., and Südhof, T. C. (2019). Synaptic neurexin-1 assembles into dynamically regulated active zone nanoclusters. Journal of Cell Biology, 218(8), 2677–2698. 10.1083/jcb.201812076

42. Ullrich, B., Ushkaryov, Y. A., and Südhof, T. C. (1995). Cartography of neurexins: More than 1000 isoforms generated by alternative splicing and expressed in distinct subsets of neurons. Neuron, 14(3), 497–507. 10.1016/0896-6273(95)90306-2

43. Wee, R. W. S., and MacAskill, A. F. (2020). Biased Connectivity of Brain-wide Inputs to Ventral Subiculum Output Neurons. Cell Reports, 30(11), 3644–3654.e6. 10.1016/j.celrep.2020.02.093

44. Wozny, C., Maier, N., Schmitz, D., and Behr, J. (2008). Two different forms of long-term potentiation at CA1–subiculum synapses. The Journal of Physiology, 586(Pt 11), 2725–2734. 10.1113/jphysiol.2007.149203

